# A neural mechanism for conserved value computations integrating information and rewards

**DOI:** 10.1101/2022.08.14.503903

**Authors:** Ethan S. Bromberg-Martin, Yang-Yang Feng, Takaya Ogasawara, J. Kael White, Kaining Zhang, Ilya E. Monosov

## Abstract

Behavioral and economic theory dictates that we decide between options based on their values. However, humans and animals eagerly seek information about uncertain future rewards, even when this information does not provide any objective value. This implies that decisions can be made by endowing information with subjective value and integrating it with the value of extrinsic rewards, but the mechanism is unknown. Using a novel multi-attribute decision making task we found that human and monkey value judgements are regulated by strikingly conserved computational principles, including how they compute the value of information and scale it with information’s timing and ability to resolve a specific form of uncertainty. We then identified a neural substrate in a highly conserved and ancient structure, the lateral habenula (LHb). LHb neurons signal the subjective value of choice options integrating the value of information with extrinsic rewards, and LHb activity both predicts and causally influences online decisions. Key input regions to LHb provide the necessary ingredients for these computations, but do not themselves signal an integrated value signal to guide multi attribute decisions. Our data thus identifies neural mechanisms of the conserved computations underlying multi-attribute, value-based decisions to seek information about the future.

## Introduction

How much would you be willing to pay to learn what your future holds? Every day we make decisions to gain concrete, physical rewards like food, water, or money, but also to satisfy more abstract desires, like our curiosity to gather knowledge about the future. How the brain makes these multi-attribute decisions, integrating the many features of extrinsic rewards and of internal abstract representations into a common currency to guide our behavior, remains unclear.

In particular, behavioral science, psychology, and economic theory propose that we choose through a process of “value-based decision making”, in which we integrate the many attributes of each option together to compute its total subjective value which then guides our choices (*1*). However, our everyday decisions often require evaluating options that have two quite different types of attributes, derived from the external world and from internal computations. Many attributes provide *extrinsic* outcomes, such as food, water, or money, or determine their statistical features such as their timing or variability. These are straightforward to study because it is possible to measure them, estimate their objective value to the organism, and compare it to the subjective value the organism assigns to it (*2–4*). Yet other attributes provide *intrinsic* outcomes, often called abstract, cognitive, or *non-instrumental* because they provide no apparent objective benefit to the organism and may not be physically measurable, yet still strongly motivate behavior (*5–7*). Thus, a critical question is how brains compute the subjective value of non-instrumental choice attributes, and how they integrate it with the value of extrinsic rewards to guide multi-attribute decisions where both must be weighed and traded off against each other.

An especially striking form of non-instrumental preference is our desire for information about uncertain future events (*8–14*). This curiosity-like information seeking is not unique to humans; it also occurs in animals including monkeys, rats, and pigeons (*15–19*). Remarkably, both humans and animals will persistently seek information about future rewards, and even pay for it, even when this information has no objective value because there is no way to use it to influence the outcome (*13, 14, 20-24*). This suggests that organisms assign information subjective value in its own right, effectively treating information itself as a form of reward. However, despite an explosion of research on information seeking in recent years in diverse fields including economics, artificial intelligence, psychology, cognitive science, and neuroscience, we are still only beginning to understand how the brain computes the value of information and uses it to guide decisions (*19, 25–31*).

Here we address two fundamental questions. First, what common principles, if any, do humans and other animal species use to compute the subjective value of information about future outcomes and integrate it with the value of extrinsic rewards? This fundamental question has been remarkably unexplored, as studies of information seeking have almost exclusively examined a single species at a time (*7*). We address this by developing a novel multi-attribute information choice paradigm for both humans and monkeys in which they choose between pairs of offers that provide different probability distributions of reward outcomes (money for humans, juice for monkeys), and different access to advance information about the outcome, allowing us to measure the subjective value of information. We found that human and monkey value judgements are regulated by strikingly conserved computational principles, including how they scale the value of information with the time the information will arrive and with its ability to resolve a specific form of uncertainty.

Second, what neuronal systems in the brain implement these conserved principles to compute the value of information, the value of extrinsic reward, and the total value of each choice alternative to drive decisions? Recent work has identified two interconnected neuronal networks with information-related activity that are prime candidates for these roles (*7*): a cortico-basal ganglia information prediction network, that contains neurons that predict information delivery and regulates information-seeking gaze shifts (*24, 32*); and the reward prediction error (RPE) network, whose signals reflect preferences for both information and primary reward, and potently regulates reinforcement learning (*18, 33–35*). However, it is unknown whether the information-related activity in these networks actually tracks the subjective value of information, and whether it has a causal role in translating that value into information seeking decisions. Indeed, previous work attempting to test this in a connected region of prefrontal cortex found neurons that did not integrate information and reward into total subjective value, instead encoding those attributes with distinct, orthogonal codes (*22*).

Here we identify a neural substrate of multi-attribute decisions at a junction point of these two networks, in an ancient epithalamic structure, the lateral habenula (LHb). Many LHb neurons tracked the full integration of information and reward into a common currency of economic value. This value signal directly regulated decisions: trial-to-trial fluctuations in LHb value signals predicted upcoming choices, while injecting weak electrical current into the LHb causally perturbed upcoming choices in a manner consistent with subtracting value from the offer. By contrast, neurons in the anterior/ventral pallidum (Pal), a part of the information prediction network and a major input to LHb, tracked the necessary attributes to compute value but only encoded them in a partially integrated manner. In particular, LHb and Pal were remarkably similar in their encoding of every individual information-related attributes, but were strikingly distinct in how they integrated them with extrinsic reward to track subjective value. Thus, our work identifies the LHb as a key substrate for conserved computations integrating information and reward into the subjective value that drives multi-attribute economic decisions.

### Conserved information valuation in humans, monkeys, and Pal and LHb neurons

We assessed how humans assign value to information in a multi-attribute information choice task (Fig 1A).). Participants (n=565) chose between a pair of offers on each trial with a mouse click. When chosen, each offer provided a monetary reward that was randomly drawn from a set of four possible outcomes, depicted as four stacks of coins (where each coin was worth 1¢; on average, participants earned a total of $9.29 from their choices in the task). Thus, different offers provided different probability distributions of rewards, which could have different levels of reward expectation (mean height of the stacks) and uncertainty (variability of the height of the stacks). In addition, each offer had a color which indicated whether it was an *Info* or *Noinfo* offer (Fig 1A). When chosen, Info offers provided an *informative cue* indicating which of the four possible outcomes would be delivered into the participant’s winnings on that trial, while Noinfo offers did not. Importantly, this information was non-instrumental and hence had no objective value because there was no way to use it to influence the outcome, and participants were clearly instructed that this was the case (*19*). Thus, each offer had multiple attributes, corresponding to multiple features of the reward distribution and the opportunity (or lack thereof) to gain information. To let us separate information preferences from color preferences, the mapping between color and informativeness was randomized for each participant and was reversed midway through the session.

**Figure 1.**
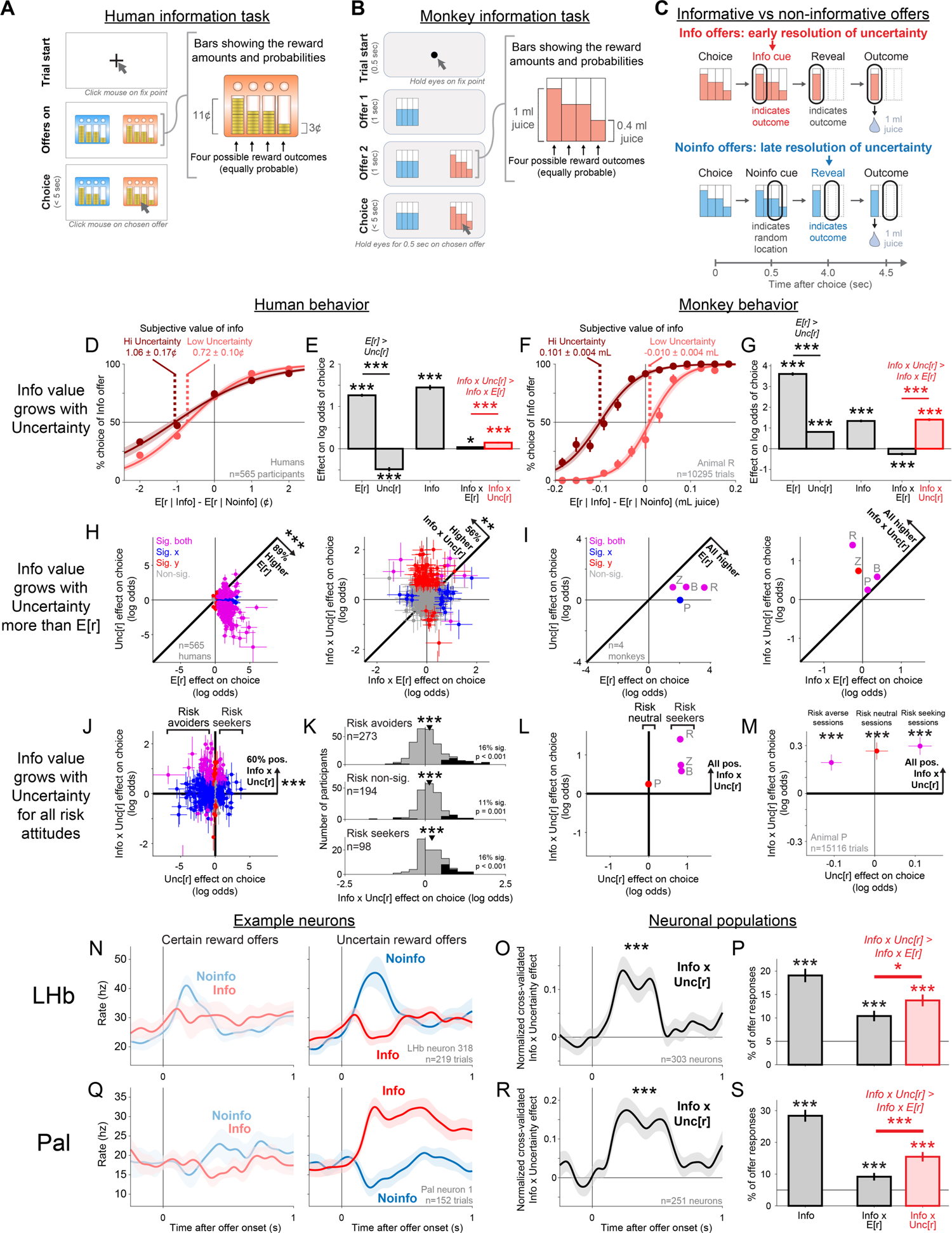
How the subjective value of information and neuronal information signals scale with uncertainty. (A,B) Choice procedure during the multi-attribute information choice tasks for humans and monkeys. (C) Info offers granted access to early information about the upcoming reward outcome, while Noinfo offers did not. (D) Psychometric curve measuring the subjective value of information based on the choice of Info vs. Noinfo offers (y-axis) as a function of the difference in their expected reward (x-axis), separately for trials where both offers had high or low reward uncertainty (dark or light red), computed using all n=565 human participants. Error bars are SE. Curve is the best fitting logistic function, shaded area is its bootstrap SE (n=200 bootstraps). Text indicates the subjective value of information implied by the curve’s indifference point, and its bootstrap SE. (E) Mean fitted GLM weights of offer attributes. Error bars are ±1 SE, *, **, *** indicate p < 0.05, 0.01, 0.001 (signed-rank test). (F,G) Same as A,B, for Animal R. (H) Humans generally placed higher value on expected reward than uncertainty (*left*, weight of E[r] vs Unc[r]), but increased the value of information more with uncertainty than expected reward (*right*, weight of Info x E[r] vs Info x Unc[r]). Each data point is one individual. Error bars are ±1 SE. Colors indicate that neither (gray), the x-coordinate (blue), the y-coordinate (red), or both (purple) are significant (p < 0.05). Text indicates the fraction of individuals above or below the identity line and its significance (binomial test). For visual clarity, axes here exclude one extreme outlier (error bar visible at the bottom of H, J). (I) Same as H, for all animals. Gray text indicates which data point came from which animal. (J) The value of information grows with uncertainty regardless of individual risk attitudes toward that uncertainty, indicated by the presence of positive weights of Info x Unc[r] (y-axis) for humans with either negative, non-significant, or positive weights of Unc[r] (x-axis). Text indicates the fraction of individuals above y=0 and its significance (binomial test). (K) Histograms of fitted Info x Unc[r] weights for individuals classified as risk avoiders, risk non-significant, or risk seekers. Black indicates individuals with significant Info x Unc[r] weights. Text indicates the fraction of individuals with significant positive weights and the p-value of whether this fraction is significantly greater than chance (binomial test). (L) Same as J, for all animals. (M) The value of information grew with uncertainty consistently in Animal P (Info x Unc[r] weight, y-axis), separately for the subsets of sessions when the animal tended to be risk averse, neutral, or seeking (Unc[r] weight, x-axis). (N) An example LHb neuron with a strongly information-related offer response that is much stronger for offers with high reward uncertainty (right) than offers with certain reward (left). (O) Mean Info x Unc[r] effect, measured as the cross-validated difference in normalized activity between Info and Noinfo offers with uncertain rewards, minus the analogous difference for certain rewards. This analysis uses all n=303 attribute-responsive neurons. Shaded area is ±1 SE. *** indicates p < 0.001 (signed-rank test). (P) Percentage of all LHb offer responses (n= 375 total neurons x 2 offers) with significant GLM weights of Info, Info x E[r], and Info x Unc[r]. Horizontal line is chance. Error bars are ±1 SE. *, **, *** are p < 0.05, 0.01, 0.001 (one-tailed binomial test). (Q,R,S) same as N,O,P for Pal neurons (n=251 attribute-responsive, n=294 total).

The information choice task for monkeys (n=4), was designed in an analogous fashion. Animals freely viewed and then chose between a pair of offers on each trial; each offer provided a reward that was randomly drawn from a set of four possible outcomes, depicted as four bars whose heights indicated their magnitudes; and offers had distinct textures indicating whether they were Info offers that provided an informative cue indicating the upcoming outcome or Noinfo offers that provided a non-informative cue. The main differences between the tasks were that monkeys were rewarded with juice rather than money; chose with eye movements rather than mouse clicks; were shown the offers sequentially (to let us measure neural responses to each offer); and were given their juice reward at the end of each trial (unlike humans whose monetary winnings were delivered as a lump sum after the experiment). Also, to ensure that animals had ample opportunity to physically prepare for the reward from both Info and Noinfo offers (*17, 18, 36*), animals were always shown a ‘reveal’ stimulus revealing the outcome shortly before reward delivery (Fig 1C). We confirmed that humans were information seeking in a similar version of the task where both Info and Noinfo outcomes were revealed at the end of each trial (Methods; Fig 3E, S1). In addition, we confirmed that monkeys had consistent information preferences in a version of the task where offers had additional visual attributes indicating the timing of cue and reward delivery (Fig 3, S2; the task used for neuronal recording).

Using these analogous tasks, we found analogous valuation of information by humans, monkeys, and neurons. We first assessed the fundamentals of how individuals and neurons valued Info vs Noinfo offers, then investigated the specific algorithms that individuals use to compute these values. In humans, many individuals were strongly information seeking. On average, humans chose informative over non-informative offers 69% of the time, and this preference was significant in 69% of participants. Humans placed high subjective value on information, as measured by their willingness to pay for it (Fig 1D), which on average over all trials was equal to 1.00 ± 0.19 cents – fully 17% of the mean expected reward of offers in the task. In monkeys, all monkeys significantly modulated their choices with information in this task (p < 0.0001 in each animal). For example, on average over all trials, the animal shown in Fig 1F was willing to pay 0.048 ± 0.002 mL of juice for information – fully 16% of the mean expected reward of offers in the task (Table S1). As monkeys performed the task, we recorded from neurons in LHb and Pal and examined their responses to the offers (n=2 animals, n=375 LHb, n=294 Pal). Both areas were highly sensitive to the offered opportunity to gain information, indicated by significant effects of offer informativeness (Info vs Noinfo; p < 0.05 in GLM fits to neuronal activity; Methods) on the firing rates of a large fraction of neuronal responses to offers (LHb 19.1%, Pal, 28.4%).

### Information value scales with uncertainty

Having found that humans and monkeys are both information seeking and that Pal and LHb neurons have information-related signals in this task, we set out to use it to answer our key questions. First, what principles do individuals follow when they use attributes of instrumentally valuable extrinsic rewards, like money and juice, to compute the subjective value of non-instrumental choices attributes, like information about future outcomes? To do so, we test how the value of information scales in our task with two major determinants of instrumental reward value: expected reward and reward uncertainty. Second, are these principles conserved between humans and monkeys, and are they implemented by the information-related signals in Pal and LHb?

To test this, we presented individuals with reward distributions that were either *safe*, where the outcome was entirely certain, or *risky*, where the outcome was uncertain (Fig 1D,F). We found that the value of information scaled strongly with uncertainty in both species. Humans were willing to pay an average of 1.06 cents for information about uncertain outcomes, but only 0.72 cents for information about certain outcomes (Fig 1D). In an analogous manner, all individual monkeys were willing to pay more for information about uncertain outcomes (Fig 1F, S4).

To quantify this more precisely in each individual, we fit each individual’s choice behavior with a generalized linear model with separate weights for each offer attribute and its interactions with information. This framework models the subjective value of each offer as a linear weighted combination of its attributes and interactions. We found that the human population had a strong and significantly positive Info x Uncertainty weight, indicating the information was subjectively valued more highly when it would resolve a larger amount of reward uncertainty (Fig 1E). The Info x Uncertainty weight was positive in 60% of individual humans; it was significant in 14% of individuals, which were predominantly positive (Fig 1H; 12% positive, more than expected by chance, p < 0.0001, binomial test; 2% negative, not significantly different from chance, p = 0.89). Similarly, all individual monkeys had significant positive Info x Uncertainty weights (Fig 1XG,I).

Crucially, the value of information predominantly scaled up with *uncertainty*, not simply with all attributes that individuals cared about. All monkeys and nearly all humans placed much greater weight on expected reward than uncertainty (Fig 1E,G; Fig 1H,I, dots below identity line), as expected from prior work. Yet when examining the interaction between these attributes and information, all monkeys and many humans were fitted with significantly higher weight on Info x Uncertainty than Info x E[r] (Fig 1E,G; Fig 1H,I, dots above identity line). Thus, while humans and monkeys generally placed more value on expected reward than uncertainty, they increased the value of information more with uncertainty than expected reward. Importantly, our ability to detect these analogous value computations was crucially aided by our experimental design using analogous tasks across species. This reduced the possibility that differences in the relative influences of Uncertainty and E[r] on information seeking might arise simply due to differences in task designs (as has been reported within-species in humans (*13, 14, 37-39*)). Thus, here we show for the first time that humans and monkeys performing similar tasks can value information in similar ways.

We next asked how these attitudes towards information about risky outcomes are related to attitudes toward risk itself, a topic with a long history in economic theory (*8, 40–43*) and psychological and computational models of information seeking behavior (*11, 23, 44-47*). According to one set of theories, the value of information derives from nonlinear weighting of information about future desirable vs undesirable outcomes, similar to how risk attitudes are theorized to derive from nonlinear weighting of the desirable vs undesirable outcomes themselves. If a single underlying mechanism regulates both phenomena, then information preferences should be tied to risk preferences (*11, 18, 41, 46*). According to other theories, information to resolve uncertainty is valued independently as an incentive in its own right, and hence may be given positive subjective value regardless of whether an individual is risk seeking or risk averse (*7, 23, 42, 44, 48*). Our data supports the latter theories, in both across-species and within-species comparisons. Across-species, consistent with previous work, humans tend to be risk averse in our task (negative weight of Uncertainty, Fig 1J; (*49*)) and monkeys tend to be risk seeking (positive weight of Uncertainty, Fig 1L; (*50*)). Yet despite this, both the human population and all monkeys were fit with significant positive weights of Info x Uncertainty, indicating that they placed higher value on information about risky, uncertain offers (Fig 1J,L). Within humans, the Info x Uncertainty weight was significantly positive in the subsets of humans with each possible risk attitude: risk-averse, risk-seeking, and risk-nonsignificant (Fig 1J,K). Similarly, within monkeys, it was significantly positive in all animals regardless whether they were risk-seeking or risk-neutral (Fig 1K), and was significantly positive even in the subsets of sessions when the one risk-neutral animal had trends for each possible risk attitude (Animal P, Fig 1M). Thus, our data is consistent with humans and monkeys treating information that resolves uncertainty about future outcomes as a separate incentive in its own right, remarkably distinct from their attitudes toward uncertainty about future outcomes itself.

We next asked whether the specific scaling of information value with uncertainty could be implemented by Pal and LHb. Indeed, the information-related activity of many neurons in these areas was significantly higher for uncertain than certain rewards (Fig 1N,O,Q,R). To quantify this, we fit each neuron’s activity with an analogous GLM to the one we used to model behavior. This revealed significant Info x Uncertainty weights in many Pal and LHb neurons (Fig 1P,S). Furthermore, more neural offer responses had significant Info x Uncertainty weights than Info x E[r] weights (Fig 1P,S, LHb p = 0.041, Pal p = 0.0007, signed-rank test). Thus, just as the subjective value of information increased more with uncertainty than expected reward, so did Pal and LHb information signals.

### Information value scales with a specific form of uncertainty

To understand why and how the value of information scales with uncertainty, is it fundamental to uncover the specific *form* of uncertainty that governs information seeking. Many distinct mathematical forms of uncertainty have been proposed to influence cognition and behavior. However, most neuroscientific studies have manipulated uncertainty using single parameters of probability distributions which make it difficult to differentiate these proposals (Fig S3). Here, we take advantage of the fact that our task can present full probability distributions of rewards to test between three hypothesized families of uncertainty measures (Fig 2A,B). One family of uncertainty measures depends only on the probabilities of the potential outcomes, most prominently Shannon entropy (*51*) and related quantities such as Kullback-Leibler divergence (*52*), which have been used to model information preferences (*44, 53, 54*) and many other forms of motivation, cognition, and neural computation (*55–59*). A second family depends only on the magnitudes of potential outcomes, including the range, which has been proposed to regulate the dynamic range of neural activity (*60–63*). Finally, a third family computes uncertainty using both probabilities and magnitudes of outcomes, most prominently including standard deviation (SD) and variance, which have been proposed to regulate risk- and information-related behavior and neural activity (*23, 50, 64-66*).

**Figure 2.**
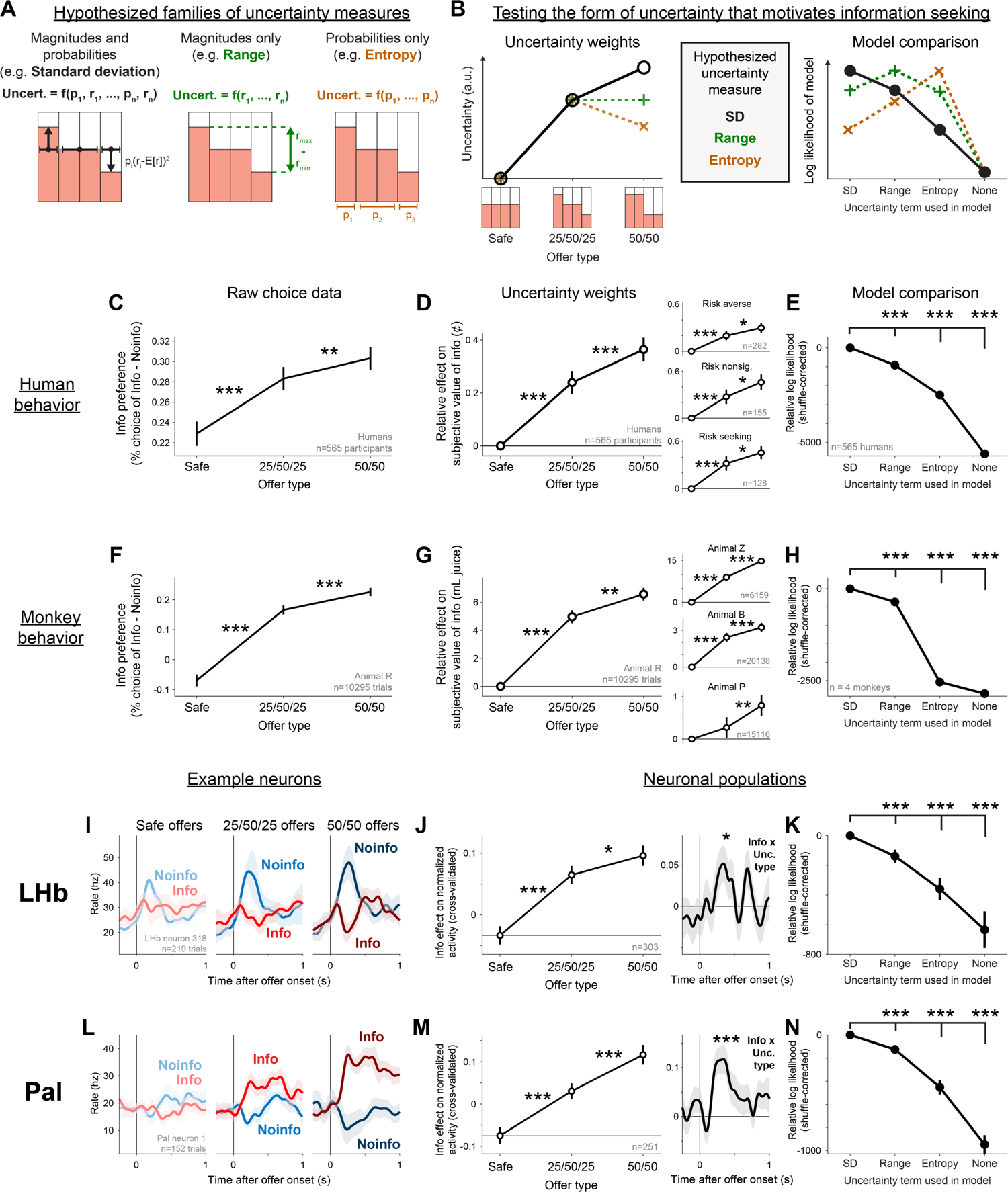
A conserved form of uncertainty scales information value. (A) Uncertainty has been hypothesized to influence behavior through different families of uncertainty measures, computed using both reward magnitudes and probabilities (e.g. SD), magnitudes only (e.g. Range), or probabilities only (e.g. Entropy). (B) We presented individuals with three offer types (Safe, 25/50/25, 50/50) designed to dissociate these families of uncertainty measures, both in terms of their effects on the subjective value of information (left) and their quality of fitting the data (right). Plots show hypothesized results according to each family of uncertainty measures. (C-E) Human information seeking is motivated by an SD-like form of uncertainty. (C) Mean difference in percentage choice of Info offers vs Noinfo offers, separately for each of the three offer types, over all n=565 human participants. Error bars are ±1 SE. *, **, *** indicate p < 0.05, 0.01, 0.001 (signed-rank tests). (D) Mean fitted GLM weights representing the effect of each uncertainty type on the subjective value of information. Insets show similar results separately for the subsets of participants classified as risk averse, risk non-significant, or risk-seeking. For this plot, to ensure this analysis is not biased to favor any of the three families of uncertainty measures, participants were classified as risk averse/seeking if they had a significant negative/positive Unc[r] effect according to any of the three models (SD model, Range model, or Entropy model; Table S2). (E) Model comparison in terms of shuffle-corrected log likelihoods relative to the SD model shows a significantly better fit for the SD model than all of the models using other uncertainty terms and the None model without an uncertainty term. Error bars are ±1 bootstrap SE (n=2000 bootstraps). *, **, *** indicate the 95%, 99%, 99.9% bootstrap CI excluded 0. (F-H) Same as C-E, but for animals, showing very similar results. F,G show data from Animal R; insets in G show data from each other animal. (I-K) LHb neuron information-related activity scales with an SD-like form of uncertainty. (I) The same example LHb neuron from Fig 1 but now separating its responses to uncertain reward offers based on the offer type, revealing that this neuron tended to have stronger information-related activity for 50/50 offers. (J) *Left:* Mean cross-validated GLM weights from fits to all attribute-responsive LHb neurons are consistent with an SD-like form of uncertainty. Error bars are ±1 SE. *, **, *** indicate p < 0.05, 0.01, 0.001 (signed-rank tests). *Right:* Mean time course of the cross-validated Info x Uncertainty Type effect on neuronal activity, measuring the enhancement of information-related activity by 50/50 relative to 25/50/25 offers (Methods). Same format as Fig 1O,R. (K) Model comparison indicates LHb neurons are significantly better fit by using an SD-like form of uncertainty. Same format as E. (L-N) Same as I-K, for Pal neurons. Model comparisons were done using all neurons where the model converged to a stable fit in all bootstrap samples (n=373 LHb, n=293 Pal).

To dissociate these families of uncertainty measures, we used two types of uncertain reward distributions: 50/50 offers where big or small rewards each occurred 50% of the time, and 25/50/25 offers where big or small rewards each occurred 25% of the time and intermediate rewards occurred the remaining 50% of the time (Fig 2B). Uncertainty measures that depend only on probabilities, like entropy, are highest for the latter since it has the most evenly spread probabilities; measures like range are indifferent, because both have the same extreme big and small reward magnitudes; and measures like SD are highest for the former, since it has the highest probability of large deviations from the mean magnitude.

Both humans and monkeys valued information based on an uncertainty measure more closely resembling SD than either range or entropy, indicated by raw choice percentages (Fig 2C,F) and fits to human and monkey behavior with separate Info x Uncertainty weights for each of the two types of uncertain offers (Fig 2D,G). This was the case for humans with all risk attitudes, and for each individual monkey (Fig 2D,G, insets). To further quantify this, we did a formal model comparison between models of behavior in which the uncertainty measure was SD, range, or entropy, and found that SD was highly significantly favored in both species (Fig 2E,H). This was the case for LHb and Pal information signals as well. The same example neurons from Fig 1 had much stronger signals differentiating Info vs Noinfo for offers with 50/50 distributions than 25/50/25 distributions (Fig 2I,L). Similarly, the population average activity had a clear Info x Uncertainty Type interaction (Fig 2J,M), and a formal model comparison indicated that neural activity in both areas was significantly better fit as tracking SD rather than range or entropy (Fig 2K,N). Thus, Pal and LHb information signals tracked the same form of uncertainty that governed information’s value in multi attribute decision making of monkeys and humans.

### Information value scales with time

We next asked whether the value of information scales with time. Information preferences have been shown to be influenced by the timing of task events (*11, 17, 18, 36, 67*), but we are only beginning to understand how these translate into subjective value and this value is computed by neural circuits (*68*). Notably, whereas standard temporal discounting scenarios only require evaluating a single event, the time of the extrinsic reward outcome (Tout), information seeking also requires evaluating the time when information will arrive – either the time of the cue (Tcue) for Info offers, or the time of the final reveal (Trev) for Noinfo offers (Fig 3A). We hypothesized that these are crucial ingredients for information valuation. Specifically, the difference between them – the time the cue comes in advance (Tadvance) – may be valued because it quantifies the temporal advantage in information (that is, how much earlier an individual will get information by choosing the informative option).

To test this in monkeys, we modified the task to augment each offer with a visual ‘clock’ (Fig 3A, n=3 animals), with three segments indicating the time durations between choice and cue; between cue and the reveal stimulus; and between the reveal and the end of the trial. After an offer was chosen, an animated ‘hand of the clock’ appeared that moved across the clock and triggered the cue, reveal, and end of the trial at the appropriate times (Fig S2). After monkeys learned the task, they retained the positive Info x Uncertainty effects shown before (Fig S4), while additionally showing strong time preferences. As expected, animals showed strong conventional temporal discounting, preferring early over late Tout (Fig 3B).

Crucially, time also strongly influenced information preferences, in a manner that depended on Tadvance. That is, animals had modest information preferences when Tadvance was short, but strong information preferences when Tadvance was long (Fig 3C). Importantly, the information preference was strongly influenced by Tcue and not simply by Tout. This could be seen clearly by the strong increase in information preference when Tadvance was manipulated by shortening Tcue while holding Tout constant (Fig 3C, right). To measure this in each animal, we fitted each animal’s behavior with a model with separate terms for the the relative subjective value of information for offers where Tcue and Tout were either early or late (Fig 3D, Methods). In all animals, the fitted value of information was significantly highest when Tcue was early and Tout was late – in other words, when Tadvance was longest (Fig 3C, all p < 0.001).

To quantify how individuals computed the subjective value of offers based on timing and all other attributes, we selected the attributes that were required to account for 99.9% of the above-chance cross-validated log likelihood of the choice data for all animals (Methods). This identified 10 value-related attributes, which we then used to model both behavior and neuronal activity. Of the 10 attributes, 5 were related to time (Methods). Two were related to the timing and amount of juice delivery (Tout and Tout x E[r]), while three were related to the interaction between timing and information (Info x Tout, Info x Tadvance, and Info x Tadvance x Uncertainty). A model comparison confirmed that these Info x Time interactions improved the fit to animal behavior (Fig 3F). Notably, the fitted weight of Info x Tadvance was significantly positive for all animals, and Info x Tadvance x Uncertainty was significantly positive in two of three animals. Thus, the subjective value of information scaled with the time it would arrive advance of the outcome, especially when there was a large amount of uncertainty for it to resolve.

To test if the value of information also scaled with time in humans in this setting (*11, 67*), we modified the human task to offer a choice between early vs. late access to informative cues (Fig 3E, n=210; Fig S1). The human population was fitted with a significant mean positive subjective value of obtaining early vs. late information (Fig 3E, p < 0.001, signed-rank test), with significant positive values occurring in a substantial number of individuals as well (21%; significantly above chance, p < 0.0001, binomial test).

Again, Pal and LHb information signals contained all the necessary components to implement these value computations in multi-attribute decision making (Fig 3G-N). Both areas had significant proportions of neurons with Info by time interactions during offer presentation (Fig 3H,L), including each of the three different interactions identified from the behavioral model (Fig 3I,M). For example, both areas contained neurons whose information signals were stronger when Tadvance was long. And just as Info x Time interactions improved the fit to behavior, they also improved the fit to neuronal information signals (Fig 3J,N).

**Figure 3.**
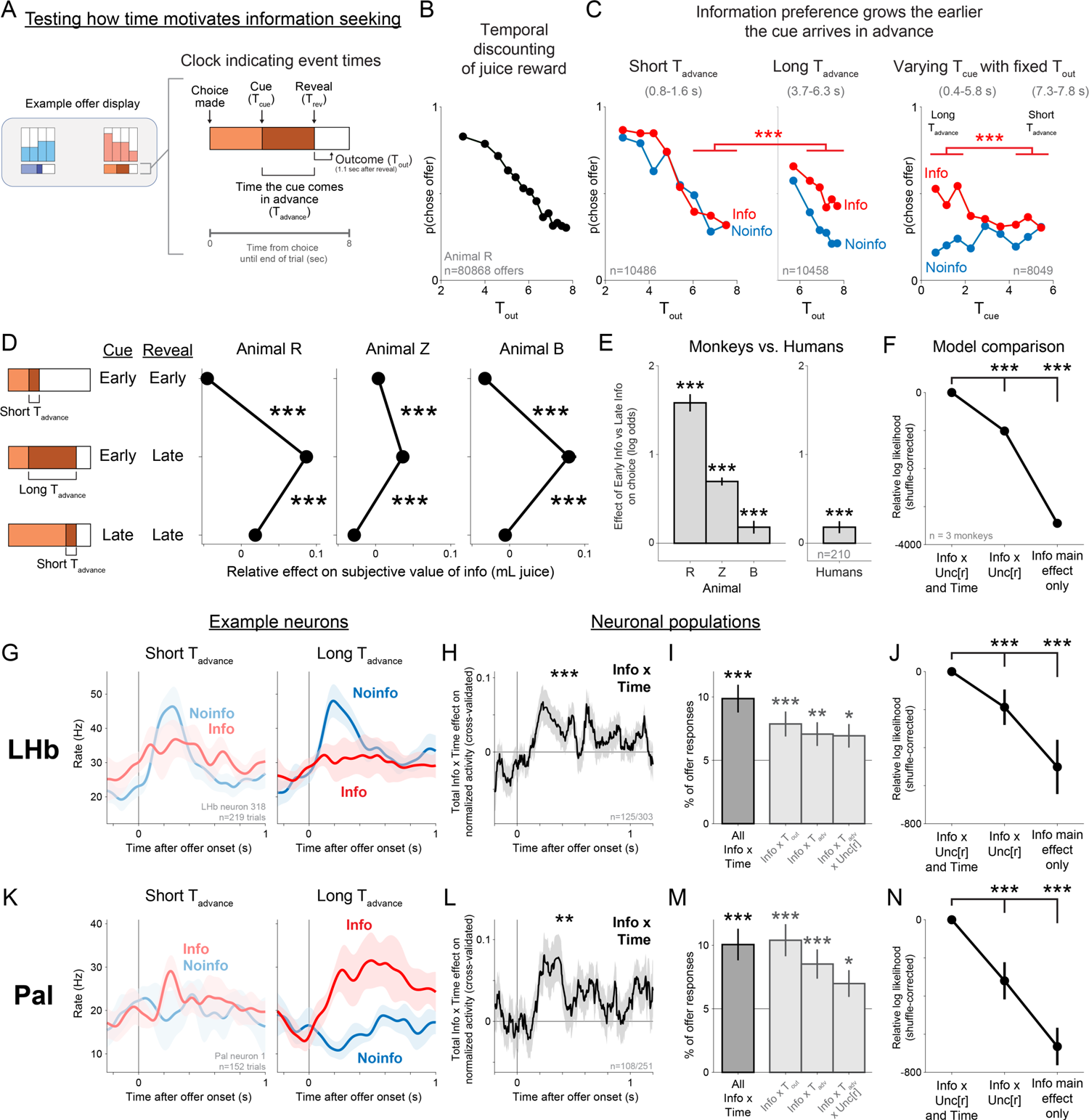
How information value scales with time. (A) We tested how information value scales with time by adding a ‘clock’ stimulus to each offer indicating the timing of cues (Tcue) and reveals (Trev), and hence their difference, the time the cue comes in advance (Tadvance). (B) We replicate conventional temporal discounting of delayed juice rewards. Plotted is the choice probably of offers as a function of their outcome delivery time (Tout) for Animal R, showing a strong preference for early over late rewards. Error bars are ±1 bootstrap SE (too small to see). (C) Information preference grows the earlier the cue arrives in advance. *Left*: same format as B, but split into Info and Noinfo offers and only plotting offers with short Tadvance, showing very weak preference for information. *Middle*: same, only plotting offers with long Tadvance, showing greatly increased preference for information. *** indicates that offers with long Tadvance had a greater difference in choice probability between Info and Noinfo offers than offers with short Tadvance (the 99.9% bootstrap CI excluded 0), when considering the same range of Tout (6-8 s). *Right*: same format, but plotting choice probability as a function of Tcue while only showing offers with Tout fixed in a narrow range (7.3-7.8 s). The preference for Info over Noinfo offers is very large when Tcue is early and hence Tadvance is long, and much smaller when Tcue is late and hence Tadvance is short. (D) Fitted relative effect on the subjective value of info of three types of offers defined by their Tcue and Trev. Crucially, information value could not be explained by either Tcue or Trev alone. Instead, all three animals tested in the task placed significantly higher value on Info when offers had early Tcue and late Trev, i.e. the longest Tadvance. Error bars are ±1 SE, *** indicates p < 0.001. (E) Cross-species comparison of the value of early vs late info. Shown is the mean fitted effect of Early vs Late Info on choice for each animal (*left*, Methods) and for the human population tested on a version of the human task that manipulated information timing (*right*, Methods). *** indicates the 99.9% bootstrap CI excludes 0. (F) Model comparison indicates that animals were significantly better fit by a model including interactions between Info and both uncertainty- and time-related variables, than by a model including interactions with uncertainty-related variables alone, or a model with no interactions. Same format as Fig 2E. (G-J) LHb neuron information-related activity is commonly modulated by information timing. (G) The same example LHb neuron from Figs 1-2, showing information-related activity when Tadvance is short vs. long. (H) The LHb population had a significant total cross-validated effect of Info x Time interaction terms (Info x Tout, Info x Tadv, Info x Tadv x Unc[r]), using the 125/303 attribute-responsive LHb cells selected by the cross-validation procedure. Shaded area is bootstrap ±1 SE. *** indicates p < 0.001 (signed-rank test). (I) Percentage of all LHb offer responses (n= 375 total neurons x 2 offers) with significant GLM weights of and Info x Tout, Info x Tadv, Info x Tadv x Unc[r]. Horizontal line is chance. Leftmost bar is the percentage with significant Info x Time effects computed by pooling the p-values of the three individual effects Fisher’s combination test (*126*). Error bars are ±1 SE. *, **, *** are p < 0.05, 0.01, 0.001 (one-tailed binomial test). (J) Model comparison. Same format as F. (K-N) Same as G-J, for Pal neurons. The population timecourse of Info x Time interactions was computed using the 108/251 attribute-responsive Pal neurons selected by the cross-validation procedure. Model comparisons were done using all neurons where the model converged to a stable fit in all bootstrap samples (n=373 LHb, n=293 Pal).

### LHb reflects integration of information and extrinsic reward into subjective value

Thus far we have shown Pal and LHb neurons encode all the offer attributes necessary to compute the subjective value of information. However, this does not necessarily mean that individual neurons encode value. Value computations can have multiple stages (Fig 4A). The many attributes of each offer must be detected, weighted, and integrated, to compute a value that is properly aligned with the individual’s preferences. Furthermore, both information- and reward-related attributes must be integrated together to compute the total subjective value of the offer to guide decisions. These stages of value computations should produce very different neural activity (Fig 4B). Neurons that are ‘labeled lines’ encoding single attributes (*69*), or that have mixed selectivity to random subsets of attributes (*70*), should generally be only weakly aligned with value. Neurons that encode a partial integration by properly weighting only a subset of attributes (*69*) should generally be partially aligned with value. Finally, neurons that encode a fully integrated value signal (*71*) should be fully aligned with subjective value (Fig 4B).

**Figure 4.**
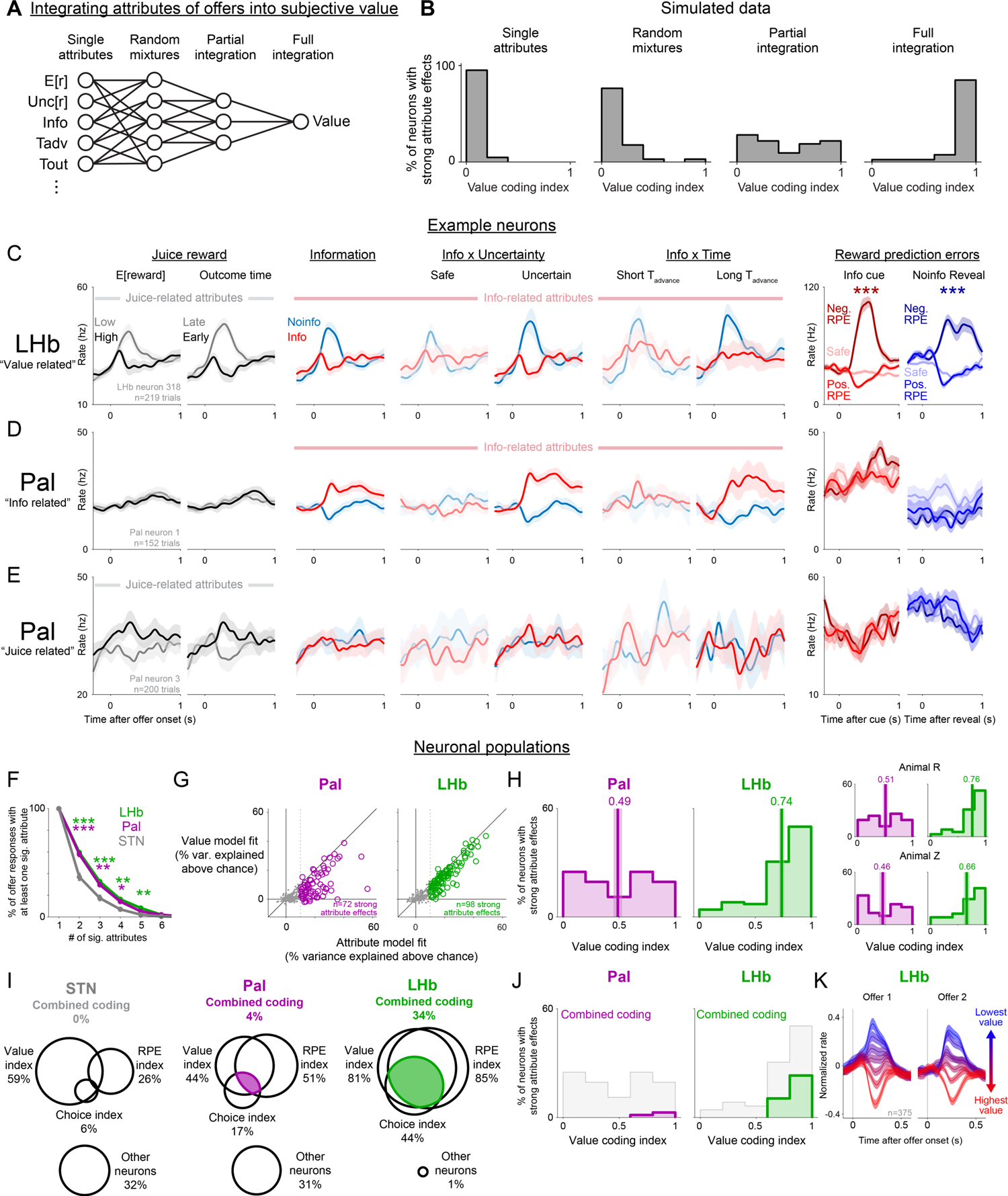
LHb neurons integrate information and reward into subjective value during multi-attribute decision making. (A) Hypotheses of neuronal attribute coding. Neurons could encode single offer attributes, random mixtures, partially integrate subsets of attributes, or fully integrate all attributes to reflect subjective value. (B) Simulations show that the Value Coding Index can distinguish full integration from the alternate hypotheses. (C) The same example LHb neuron from Figs 1-3, showing responses to attributes related to juice reward (*left*; E[r] and Tout) and information (*middle*; Info, Info x Unc[r], and Info x Tadvance; same format as Figs 1,2,3), and juice reward prediction errors (*right*; induced by informative cues on Info trials (red) and the final reveal on Noinfo trials (blue); medium, dark, and light color shades indicate positive RPEs (> 50 µL), negative RPEs (< −50 µL), and safe offers with no RPE). (D) The same example Pal neuron from Figs 1-3 is sensitive to attributes related to information (middle) but not juice reward (left) or RPEs (right). (E) A second example Pal neuron is sensitive to attributes related to juice but not information or RPEs. (F) LHb and Pal attribute-responsive neurons (green and purple) have significant effects of more attributes than attribute-responsive STN neurons (gray). *, **, *** indicates p < 0.05, 0.01, 0.001 between LHb or Pal and STN. This plot includes all offer responses that had a significant effect of at least one attribute (treating each neuron’s responses to offer 1 and offer 2 separately; totaling n=470, 387, and 186 such offer responses in LHb, Pal, and STN respectively). (G) Percent of variance in the neuronal data explained above chance for each attribute-responsive Pal neuron (left, purple) and LHb neuron (right, green), when fit using the full attribute model (x-axis) or the simple value model (y-axis). Vertical dashed line indicates the threshold for classifying cells as having strong (colored circles) or weak (gray circles) attribute effects. (H) Stronger value coding in LHb than Pal, shown by histograms of Value Coding Indexes from the neurons in each area with strong attribute effects. Text and vertical line indicates the mean, shaded area is ±1 SE. *Inset*: same separately for Animal R (top) and Z (bottom). (I) Many LHb neurons, but few Pal or STN neurons, combined offer value coding with other key forms of motivational coding. Venn diagrams show overlap among the neurons with strong attribute effects, between neurons with high Value Coding Indexes (left), significant RPE Coding Indexes (right), and significant Choice Predictive Indexes (center), as well as neurons that met none of those criteria (bottom). Colored area and text indicates the neurons with combined coding of all three properties. (J) Same as H, highlighting the subset of neurons with combined coding of all three properties. (K) LHb population activity closely reflects the value of each offer, in a negative manner with higher firing rates for lower valued offers. Shown is the population average normalized firing rate of all LHb neurons in response to Offer 1 (left) and Offer 2 (right), separately for each of 7 bins of offer subjective values (derived from the model fit to behavior).

Indeed, we found that many LHb and Pal neurons encode attributes in strikingly different ways, consistent with different stages in value computations. The LHb neuron in Fig 4C tracked a large number of attributes needed to compute the total subjective value of an offer, including attributes related to both juice reward and to information. By contrast, the Pal neurons in Fig 4D,E each strongly encode multiple attributes of each offer, but do not integrate them in a manner resembling total subjective value. The first neuron strongly activates for Info, and scales up this activation with attributes that scale up the value of info (Uncertainty and Tadvance); however, it does not track attributes related to juice (Fig 4D). The second neuron activates for attributes related to the value of juice (high E[r], early Tout), but has little response to attributes about information (Fig 4E).

These response patterns were common in the neural populations as a whole. We first fit each neuron’s offer responses with the model described above, and found that on average, both LHb and Pal neurons were significantly sensitive to similar numbers of offer attributes (Fig 4F). We then asked whether they integrated those attributes in a manner resembling subjective value. To do this, we calculated a Value Coding Index defined as the ratio of the above-chance variance in a neuron’s activity that could be explained by a model with a single term reflecting total subjective value, versus a model with separate weights for each attribute (Value Model vs Attribute model, Fig 4G,H). Thus, an index of 1 indicates that all the above-chance variance in its attribute-related activity can be explained by subjective value, while an index of 0 indicates that attributes are encoded orthogonally to subjective value. We analyzed all neurons with strong attribute effects, meaning that the fitted attribute model explained at least 10% more response variance than expected by chance, with at least one attribute having a significant effect (Methods). The result was clear. LHb neurons predominantly had very high value coding indexes, with many very close to the maximum index of 1, consistent with full integration (Fig 4H). Pal neurons had a roughly uniform distribution of indexes, reflecting highly diverse degrees of integration, consistent with partial integration (Fig 4H). Thus, as a whole, LHb had significantly greater value coding indexes than Pal (rank-sum test, p < 0.001; Animal R, p < 0.001; Animal Z, p = 0.024). Indeed, even without selecting neurons based on their task responsiveness or response properties, the simple population average LHb firing rate closely tracked the total subjective value of each offer as it was presented (Fig 4K).

This integrated subjective value signal in LHb neurons makes them well suited for several roles in motivated behavior. First, this value signal could be present in RPE coding neurons, in which case it could be used as an RPE signal to drive reinforcement learning of value-based behavior (*34*). Second, if this value signal truly reflects the subjective value that governs choice, it could be used to monitor or drive choice behavior. We therefore tested whether LHb motivational signals are organized to support these roles, and whether this organization is specific to LHb or whether it could be inherited from inputs such as Pal. To do this, we classified LHb neurons as offer value-related if they had high Value Indexes (≥ 0.6; Methods), and then calculated two additional indexes to quantify whether the same neurons have activity related to RPEs and to choice behavior.

First, we observed that LHb offer responses resembled classic LHb negatively-signed encoding of RPEs (*34, 72*), since on average neurons were inhibited by high value offers (better than predicted) and excited by low value offers (worse than predicted). We therefore computed an RPE Index to test if these LHb neurons also encoded RPEs in response to feedback about the trial’s reward outcome (which in our task came from cues on Info trials and reveals on Noinfo trials; Methods, Fig 4C). Indeed, a large number of LHb neurons had significant RPE Indexes (Fig 4I, S5), including the example neuron which had clear RPE coding for both cues and reveals (Fig 4C).

Second, if LHb offer responses track the subjective values that govern decisions, then trial-to-trial variations in LHb value signals should predict variations in choice. Furthermore, this relationship should hold above and beyond the predictions of our neural and behavioral models (which are fitted to the pooled data over all trials, and hence cannot predict trial-to-trial variations in subjective values). For example, if a LHb neuron treats a particular offer on a particular trial as less valuable than the model predicts, then the animal should be less likely to choose that offer than the model predicts. To test this, we computed a Choice Predictive Index measuring the extent to which the neuron’s residual offer responses predicted residual choices (Methods; described in full detail in the next section and illustrated in Fig 5). Indeed, many LHb neurons had significant Choice Predictive Indexes (Fig 4I, S5), including the example neuron which strongly predicted upcoming choices (Fig 5).

**Figure 5.**
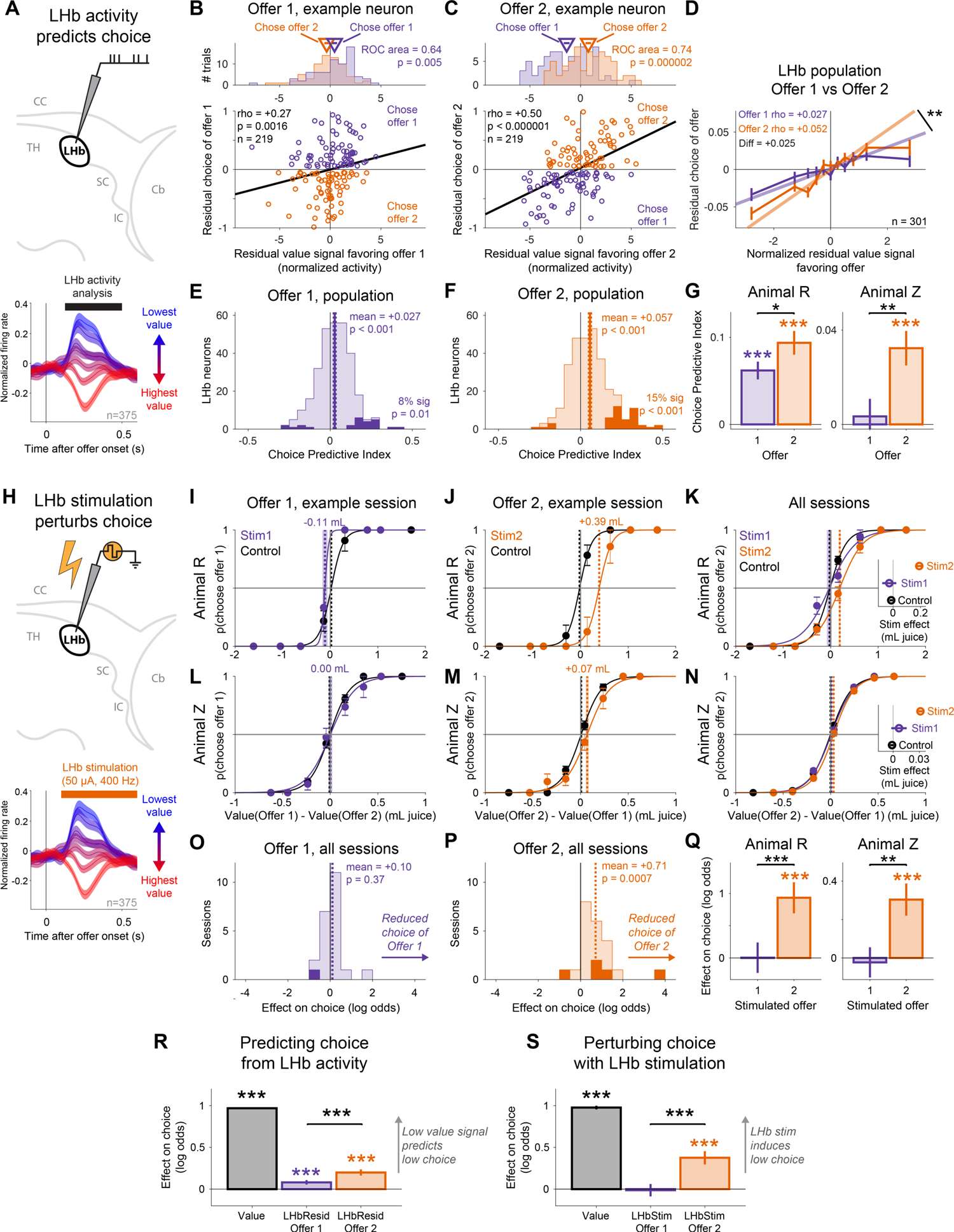
LHb both predicts and causally influences online decision making. (A) Schematic of testing whether LHb value-related activity is choice-predictive. Black bar indicates analysis time window. (B,C) The same LHb example neuron from Figs 1-4 had trial-to-trial variations in value-related activity during Offer 1 (B) and especially during Offer 2 (C) that predicted trial-to-trial variations in choice. Scatterplots show correlation between residual neuronal value signals during the response to each offer (x-axis) vs. residual choice of that offer during the subsequent decision (y-axis). Each dot is one trial. Colors indicate trials when the animal chose Offer 1 (purple) or Offer 2 (orange). Lines are the best linear fit from type 2 regression. Text indicates rank correlation and its p-value. *Top*: marginal histograms of residual neuronal value signals for choices of Offer 1 vs Offer 2; text indicates ROC area between them and its p value. (D) LHb population average relationship between normalized residual value signals favoring an offer (x-axis) and residual choice of that offer (y-axis) was significantly positive for both offers but had a significantly more positive slope for Offer 2 (** indicates the 99% bootstrap CI excluded 0). Error bars are ±1 SE. Shown are all n=301 attribute-responsive LHb neurons with sufficient data from both offers for this comparison. (E,F) Histograms showing each neuron’s Choice Predictive Indexes for Offer 1 (E) and Offer 2 (F). Dark area indicates neurons with significant indexes (p < 0.05). Dashed vertical line and text indicates mean and p value of the median being different from 0 (signed-rank test); text in lower right indicates the fraction of neurons with significant positive indexes and its p value for being greater than chance (one-tailed binomial test). (G) Both animals had significantly higher mean Choice Predictive Indexes for Offer 2 (orange) than Offer 1 (purple). *, **, *** indicates p < 0.05, 0.01, 0.001 (signed-rank tests). (H) Schematic of testing whether LHb stimulation perturbs choice. Orange bar and text indicate stimulation time window and parameters. (I,J) Psychometric curve from an example session in Animal R showing how LHb stimulation during Offer 1 (I, “Stim1”, purple) or Offer 2(J, “Stim2”, orange) affected the probability of choosing that offer as a function of the difference in subjective value of the two offers (as derived from the model fit to behavior on control trials (black), and converted to units of mL juice). Text and vertical dashed line indicates the indifference point; shaded area is ±1 SE. (K) Psychometric curves from Animal R pooling data from all sessions showing how stimulation during Offer 1 and Offer 2 influenced choice of Offer 2 relative to control trials. *Inset*: Indifference points for each condition indicate that stimulation during Offer 2 reduced choice of Offer 2. (L-N) Same as I-K, for Animal Z. (O,P) Histograms of GLM fitted effect of stimulation during Offer 1 (O) or Offer 2 (P) on choice of that offer. Positive effects indicate a reduction in the log odds of choosing the stimulated offer. Dark areas are single sessions that are significant (p < 0.05). Dashed vertical line and text indicates the mean effect and p value of the median being different from 0 (signed-rank test). (Q) Fitted main effect of stimulation on choice in each animal, pooling data over all sessions. Positive effects indicate a reduction in the log odds of choosing the stimulated offer. *, **, *** indicate p < 0.05, 0.01, 0.001. (R) Trial-to-trial variations in LHb activity aid in predicting choices. Shown are fitted weights from a GLM predicting choices based on the model-derived subjective values of the offers (Value, black) and residual normalized value signals from LHb responses to Offer 1 (purple) and Offer 2 (orange). LHb residual signals significantly predict choice, and LHb signals during Offer 2 are given significantly more weight than signals during Offer 1. Error bars are ±1 SE. *** indicates p < 0.001. (S) LHb stimulation perturbs choices. Shown are fitted weights from a GLM predicting choices based on the model-derived subjective values of the offers (Value, black) and LHb stimulation during either Offer 1 (purple) or Offer 2 (orange). Positive weights indicate a reduction in the log odds of choosing the stimulated offer. LHb stimulation during Offer 2 significantly reduces choice and does so significantly more than stimulation during Offer 1.

Using these indexes, we found that these three coding properties were partially combined in many single Pal neurons, but were fully combined in many single LHb neurons. Specifically, consistent with partial combination, both Pal and LHb had significant pairwise correlations between all three coding indexes (all p < 0.05; Fig S5). However, LHb had much higher overlap between the three coding properties – high Value Index, significant RPE Index, and significant Choice Predictive Index (Fig 4I). Notably, only LHb had a substantial proportion of “fully combined” neurons – that is, neurons that coded all three properties and did so with the same coding sign. This occurred in 34% of LHb neurons with strong attribute effects but only 4% of Pal neurons with strong attribute effects (Fig 4I,J). These “fully combined” neurons occurred much more often than chance in LHb (p < 0.001, permutation test) but not in Pal (p = 0.09), and hence to a significantly greater extent above chance in LHb than Pal (p = 0.0025).

Our multi-attribute task design was crucial to detect this difference in coding between Pal and LHb. For example, the Pal cell in Fig 4E would have appeared to encode total subjective value if we had only manipulated juice and not information. This suggests that Pal contains subpopulations of neurons that partially integrate attributes, which would be suitable to motivate specific actions (e.g. information- or juice-specific behaviors), while other Pal neurons integrate attributes more fully, suitable to regulate LHb activity and decision making. These Pal coding properties are especially notable because they are not simply found in all basal ganglia nuclei that project to LHb. To test this, we recorded n=185 neurons from the subthalamic nucleus (STN), which has strong connections with both Pal and LHb. STN had strong signals related to offer attributes (Fig 4F, S5), but compared to Pal, its neurons coded fewer attributes of offers (Fig 4F) and combined fewer coding properties (Fig 4I). Thus, our data identify Pal as a site of diverse integration of motivational attributes, and LHb as a site of predominant integration into subjective value.

### LHb value signals predict and causally influence online multi-attribute decisions

We hypothesized that LHb signals tracking subjective value have a causal role in decisions. If so, this would extend the role of LHb phasic motivational signals beyond their classic function in trial-and-error reinforcement learning to online control of sophisticated, multi-attribute decisions. As we showed above, the LHb responses to each offer in our task resembled encoding of RPEs with each offer triggering an RPE related to its subjective value, and these same neurons tended to have strong RPE signals at other times in the task when outcomes were cued and revealed (Fig 4C,I, consistent with classic findings in the LHb (*34, 72*)). However, LHb RPEs have primarily been implicated in reinforcement learning rather than online decision making (*35, 72–77*). If our hypothesis is correct, then the LHb response to each offer should predict and causally influence the animal’s decision about whether to choose that offer. Furthermore, given the negative sign of LHb value-related signals, LHb activity should be negatively associated with choice.

To test this hypothesis, we first asked whether variations in LHb value signals predict variations in choice behavior (Fig 5A). For each neuron, we computed a Choice Predictive Index as the correlation between its residual offer response and the animal’s residual choice (i.e. the correlation between the response and choice, after subtracting out the neuron’s predicted offer response and the animal’s predicted choice based on the neural and behavioral model fits; Fig 5B,C). Many neurons had strong and significant indexes (Fig 5E,F). If this LHb choice-predictive activity was related to decision making, then one could expect it to become most prominent after the animal observed both offers and could make a decision between them (*78*). To test this, we computed the Choice Predictive Index separately for the responses to the first and second offer. LHb value signals were significantly choice-predictive for both offers, but were significantly more choice predictive during offer 2 (Fig 5D-F) in both animals (Fig 5G).

To test whether the LHb causally influences online decisions, we manipulated LHb spiking activity using weak electrical stimulation at the same time LHb neurons signaled the integrated value of an offer (Fig 5H; n=2 animals; 50 µA, 400 Hz, 500 ms). On each trial, we either stimulated during the LHb response to offer 1, offer 2, or did not stimulate during either offer (Methods). If our hypothesis is correct, then adding spikes to the LHb response to an offer should cause the animal to treat it as if it had lower subjective value, and hence choose it less often. Furthermore, since LHb activity was more choice-predictive during offer 2, LHb stimulation during offer 2 might have a greater influence on choice.

Indeed, LHb stimulation substantially influenced decisions (Fig 5I-Q). This could be seen as a shift in the psychometric curves relating estimated offer values to choice, such that both animals were less likely to choose the stimulated offer, particularly when stimulation occurred during Offer 2 (Fig 5I-N). To quantify the effect of stimulation, we extended our behavioral model to include additional attributes indicating the presence of stimulation during each offer (Stim1 and Stim2). Stimulation had a significant effect on choice, and it was significantly more potent during offer 2 than offer 1 (Fig 5O,P; Stim2 p = 0.000003, Stim1 p = 0.98, Stim 2 > Stim 1, p = 0.0001). This occurred consistently across sessions (Fig 5O,P) and animals (Fig 5Q). The fitted model parameters are consistent with LHb stimulation during offer 2 leading animals to treat that offer as if it had less subjective value, equivalent to a reduction in its expected yield of juice by 0.164 mL in animal R and 0.036 mL in animal Z. Control analysis showed that the effects of LHb stimulation on choice could not be explained as simple direct effects of LHb stimulation on motor actions such as choice reports (Fig S6).

We next asked whether and how LHb activity could improve our ability to predict behavioral choices, by augmenting our behavioral models with additional terms representing either natural variations in LHb residual activity (Fig 5R), or artificial, stimulation-induced variations in LHb activity (Fig 5S). There was a strong similarity between the fitted parameters of these models, such that both natural and artificial variations in LHb activity were significantly choice predictive, and were significantly more so during offer 2 (Fig 5R,S).

Finally, given our finding of highly integrated LHb value signals, LHb stimulation should influence decisions by influencing the total, integrated value of the offer, not by selectively modulating the value of specific attributes (e.g. juice, information, etc.). Indeed, in both animals, model comparisons indicated that the influence of stimulation on decisions was largely explained by a main effect, equivalent to subtracting a fixed amount of value from the offer. There was no further significant improvement in the fit by including interactions between stimulation and offer values or specific offer attributes (Table S4).

Thus, our data identify the LHb as a key site that implements conserved information value computations by integrating information and reward into subjective value to causally influence online decisions.

## Discussion

We found that humans and monkeys value information about uncertain future outcomes and integrate it with the value of primary rewards through remarkably conserved computations, even to the extent of measuring uncertainty with the same functional form and using it to motivate information seeking in the same manner. Furthermore, we find that these conserved value computations are propagated in an evolutionarily conserved ancient epithalamic structure, the LHb. LHb contains single neurons tracking the total subjective value of offers, integrating both informational and primary reward values, and causally influences online decisions. By contrast, neurons in major basal ganglia inputs to LHb, anterior/ventral Pal and STN, commonly encoded attributes or partially integrated values but rarely reflected the integrated subjective values that drive multi attribute decisions.

The LHb is an ancient structure conserved in even the most primitive extant vertebrates (*79*). It has been consistently implicated in primary reinforcement processes across species as diverse as fish, rats, and primates (*34, 80, 81*). Our work shows that LHb also has a key role in the sophisticated motivational value computations that primates use to make multi-attribute decisions, including economic judgements trading off extrinsic primary rewards with the more abstract intrinsic reward of gaining information to resolve uncertainty about the future.

Our findings shed new light on the Pal-LHb pathway in value computations and decisions. Neurons in both areas have been reported to respond to stimulus attributes, such as expected reward, that are needed to compute subjective value (*82–85*), and to encode RPEs (*82, 86, 87*), suggesting the hypothesis that Pal neurons could motivate behavior by directly providing LHb neurons with their value and RPE-related signals. Here we show that in primates Pal does have the necessary signals to accomplish this, but primarily at the population level, not single Pal neurons. Our multi-attribute task revealed that most Pal neurons only partially integrated offer attributes, and very few of them combined strong value signals with the choice-predictive and RPE-related activity that were common in LHb neurons. Instead, our data indicates that Pal contains a diverse repertoire of neurons which resemble intermediate stages of value computations, identifying functional subsets of neurons to be further defined by anatomical and molecular studies. These Pal subpopulations could be suitable to motivate attribute-specific forms of behavior tailored to attributes of juice (e.g. thirst (*88*)), information (e.g. curiosity and gaze shifts (*32*)), or other incentives, which may explain why Pal is implicated in such diverse forms of motivated behavior (*83*). Furthermore, these subpopulations could be further integrated to compute the subjective value signals in LHb to motivate total value-guided behavior. Thus, rather than value and RPE-related activity being present in similar forms across the network, our data is consistent with distinct subcortical areas having roles in distinct steps of subjective value and RPE computations.

More precisely, our data implicates the LHb in tracking and influencing the subjective value of options that drive online decisions. While it was established early on that LHb responses are related to reward and punishment (*82, 89*), there are multiple motivational systems in the brain that track similar motivational variables for radically different functions (e.g. learning vs. online decisions, different forms of learning, and different forms of decisions (*90–94*)). The LHb has been most heavily implicated as a source of teaching signals in a reinforcement learning system including dopamine neurons and the basal ganglia (*95–97*). Yet in many contemporary theories, the “value” in these teaching signals is quite different from the subjective value that drives online decisions (*98*). Further, while LHb stimulation has been shown to strongly influence behavior, this is often a slow change over trials indicative of a learning process (*75, 99–101*), or a general aversion to stimulated contexts as in real-time place aversion (*75*). There is much less data on the role of LHb activity as decisions are made (*35, 76, 100, 102-104*). Here we demonstrate converging evidence that LHb neurons reflect value and influence online decisions: (1) the lion’s share of LHb offer responses were explained by subjective value; (2) variations in these value signals were choice-predictive; (3) LHb stimulation altered decisions as if the value of the stimulated offer was reduced; (4) both the choice-predictiveness of LHb value signals and the causal influence of LHb stimulation were greater when both offers were presented and animals could decide between them. Thus, the LHb does not simply signal RPEs for the purpose of adjusting estimated values over long timescales during reinforcement learning (e.g. across repeated presentations); these signals also directly influence ongoing decisions (Supplemental Discussion).

Our data also has important implications for a fundamental question in neuroscience: what form of uncertainty motivates behavior, and how? While it is well known that a great many aspects of cognition, motivation, and behavior are regulated by uncertainty (*105*), and many families of uncertainty measures have been proposed for this purpose, it is rarely possible to test between them with conventional task designs manipulating only a small number of outcome attributes at a time (Fig S3; Supplemental Discussion). Here we find that humans and monkeys scale up the value of information with a specific family of uncertainty measures resembling the standard deviation or variance of rewards, even for individuals with diverse attitudes toward risk *per se*. Further, Pal and LHb neurons scale their information signals in the analogous manner needed to compute the value of information. This suggests that the brain computes the value of information by tracking the full suite of reward statistics that produce uncertainty in natural environments, including both probabilities and magnitudes, and this places important constraints on the underlying neural computations (Supplemental Discussion). Further, information value computations were surprisingly robust, not only generalizing across species but also across very different behavioral regimes. Humans learned the task by written instructions, and chose with mouse clicks for monetary rewards in a single session sitting at their own computer; while monkeys learned across many sessions, and chose with eye movements for juice rewards in an experimental booth. Thus, the conserved computations humans and monkeys used to endow information with value are not artifacts of a specific behavioral regime, and are likely to generalize across a range of natural and experimental settings.

Finally, our work shows that a neuroscience approach using both humans and monkeys can identify conserved computations underlying complex multi-attribute decision making and tie them to specific neuronal substrates. This approach will be necessary to uncover the neuronal mechanisms underlying the complex decisions that we and other primates make in everyday life, which often involve making tradeoffs between multiple potential goals, which we can prefer for both instrumental and non-instrumental reasons – for example to gain reward and information for curiosity’s sake. Further, just as work in both humans and monkeys has been crucial for understanding movement disorders and developing neuroscientific treatments (*106*), the same is likely to be true for disorders of mood and cognition that impair complex decision making and often include maladaptive information seeking strategies (*107, 108*).

The LHb, in particular, has been heavily implicated in psychiatric disorders including depression and schizophrenia, as well as substance abuse, and has been identified as a major influence on models of depression and drug use in animals (*109–111*). Interventions targeting LHb are being evaluated to treat treatment-resistant depression and other mental disorders in humans (*112, 113*). Yet how alterations in LHb and its input structures produce changes in mood and decisions in everyday life is still unclear. Our work suggests a potential explanation for why alternations in LHb function can have such broad effects on mood and motivation and everyday life, as the highly integrated nature of the LHb neural code means that its subjective value signals could be engaged in a wide variety of environments with diverse motivational goals. Further, the integration of information and reward into an immediate impact on decisions means that LHb can fulfill two complementary functions simultaneously: (1) a teaching signal to learn which environments produce valuable outcomes; and (2) an immediate motivational push to enter environments that will provide both more reward, *and* more informative feedback about rewards to obtain a higher quality teaching signal.

Indeed, several forms of information seeking are altered in individuals with traits or disorders affecting mood and cognition (*114–116*), including diseases or impairments of neuromodulators that LHb potently regulates, including the serotonin and dopamine systems (*117–120*). Thus, our work raises the possibility that disordered LHb signals produce mood alterations not only due to impairing an individual’s motivation for extrinsic rewards, but also by sapping their motivation to seek information from their environment to learn about the rewards that are available in the world around them.

## Author contributions

ESBM designed the monkey task. YYF, ESBM, and IEM designed the human task. ESBM, YYF, TO, and JKW performed monkey behavioral and neural experiments. KZ performed monkey behavioral experiments. YYF performed the human experiments and monkey stimulation experiments. ESBM and YYF analyzed the data. ESBM wrote the paper in consultation with YYF and IEM. IEM guided the research and secured the funding.

## Acknowledgements

We thank Ms. Kim Kocher for fantastic animal care and training, Dr Okihide Hikosaka for many helpful discussions during the initial development of the monkey task, and Dr Lawrence Snyder for help with brain imaging. This work was supported by the National Institute of Mental Health under the Conte Center Grant on the Neurocircuitry of OCD MH106435, and R01MH110594 and R01MH116937 to IEM.

## Methods

### General procedures

A total of 824 human participants completed tasks using Amazon’s Mechanical Turk service. All participants provided informed consent (mean age = 36.06 years, SD of age = 11.15 years; 410 females). Participants were monetarily compensated based on their performance, as detailed below. The study was approved by the Washington University Institutional Review Board. Four adult male rhesus monkeys (*Macaca mulatta*) participated in experiments (Animals R, Z, B, and P). All animal procedures conformed to the Guide for the Care and Use of Laboratory Animals and were approved by the Washington University Institutional Animal Care and Use Committee.

### Behavioral task for humans

The main multi-attribute information choice task for humans involved making choices between offers which provided opportunities to earn virtual coins representing monetary rewards (n = 580 participants; mean age = 36.20 years, SD of age = 11.06 years; 294 females). We analyzed the data from all participants whose behavior contained enough variability for the generalized linear model described below to converge to a valid fit, e.g. participants would not be included if they exclusively followed a simple deterministic strategy such as always choosing the offer on the right side of the screen (though we observed a very similar pattern of results when analyzing all data without this restriction). This produced a dataset of n = 565. Participants were paid $0.01 for each coin they earned in the task. Participants learned the task by reading written instructions at their own pace. The instructions explained the task structure, the different types of offers and their attributes, and the rewards. After reading instructions, participants completed 10 practice trials before beginning the actual task.

The task consisted of 3 blocks of 50 trials each, for a total of 150 trials. Participants had 45 minutes to complete all trials. In addition, participants had a total of 10 minutes of break time that they could take as they wished between blocks. At the beginning of each trial a central cross appeared on the screen which the participant had to click to initiate the trial. Once the cross was clicked, two offers appeared on the screen. The participant had 5 seconds to choose an offer by clicking on it. If a choice was not made in time, an offer was chosen randomly by the computer. Once a choice was made, the unchosen offer disappeared while the chosen offer displayed an animation sequence which lasted ∼3.7 seconds (Fig S1). At the end of the animation, the participant had to click the offer to advance to the next trial.

Each offer featured a reward distribution that was visually depicted with 4 stacks of coins. If an offer was chosen, one of its 4 stacks would be selected uniformly at random as the outcome, and the number of coins in that outcome stack would be added to the participant’s total winnings. Each reward distribution was defined by an expected reward and type of uncertainty. Expected reward could be 5, 6, or 7 coins. Type of uncertainty could be Safe (all 4 stacks of coins were the same height), 25/50/25 (one stack was 4 coins less than the mean, two stacks were the mean, and one stack was 4 coins more than the mean), or 50/50 (two stacks were 4 coins less than the mean and two stacks were 4 coins more than the mean). Each offer also could be informative or non-informative, indicated by its color (blue or orange). If an informative offer was chosen, the animation sequence ended by highlighting the outcome stack and displaying a text message that informed the participant how many coins they had earned on that trial (e.g. “You won 7 coins!”). If a non-informative offer was chosen, the animation sequence ended by highlighting none of the stacks and displaying a text message that did not inform the participant (“You woncoins!”). The initial mapping between color and informativeness was randomized for each participant, and was reversed approximately halfway through the task (block 2, trial 20). The instructions informed participants that this reversal would occur at some point during the task.

All offer pairs were generated randomly on each trial for each participant. On 75% of trials, each offer’s features (expected reward, uncertainty type, and informativeness) were selected independently and uniformly at random across each feature’s possible values. On the remaining 25% of trials, offer features were selected such that one offer was informative and the other was non-informative, but both offers had the same expected reward and uncertainty type (selected independently and uniformly at random). For this study we combined data from three slightly different versions of the task which produced similar results. In the first version, 25/50/25 and 50/50 offers were referred to by the instructions as “moderate risk” and “high risk” respectively; in the other versions they were both referred to as “risky” offers. In the first two versions, on the 25% of trials where offers were generated to have the same reward distribution but different informativeness, the informative offer was placed on the right; in the other version, the offer positions were randomized. To test how participants understood the mapping between colors and informativeness, we included additional “question trials” (2 per block, after trials 30 and 50). On these question trials, participants were shown pictures of two offers with empty coin stacks, one informative and one non-informative. On half of question trials they were asked to click on the informative offer. On the other half they were asked to click on the non-informative offer. There was no time limit to respond, although task time continued to count down. Each correct response earned the participant $0.10. The great majority of participants answered most question trials successfully, so we included all participants in our analysis; restricting analysis based on question trial performance produced a similar pattern of results.

A second task for humans manipulated information timing (n = 244 participants, 210 with valid model fits; mean age = 35.73 years, SD of age = 11.37 years; 116 females; no overlap of participants between the two tasks). The task was similar to the main task described above, with the following differences (Fig S1). The task consisted of 2 blocks of 50 trials each, for a total of 100 trials. The uncertainty type for all offers was 50/50. The animation after choosing an offer was longer (∼14 seconds). Instead of the offer colors indicating informative vs. non-informative offers, they indicated early-information vs. late-information offers. If an early-information offer was chosen, the outcome stack was highlighted as soon as the animation began, immediately revealing the outcome. If a late-information offer was chosen, the outcome stack was highlighted at the end of the animation, revealing the outcome only at the end of the ∼12 second delay. In either case, at the end of the animation a text message appeared informing the participant how many coins they earned on that trial (e.g. “You won 7 coins!”). Of the 25% of trials in where the offers were generated to have the same offer distribution but different informativeness, n=24 were generated deterministically (8 for each of the 3 possible expected reward values) and n=1 was generated by choosing its expected reward value uniformly at random. Question trials were presented after trials 10 and 40 in each block.

### Behavioral task for monkeys

The multi-attribute information choice task for monkeys had two versions. The first version involved making choices between offers for juice rewards with fixed event timing (animals R, Z, B, and P). The second version of the task augmented the offers to allow different event timings (animals R, Z, and B). Recording and stimulation experiments were then carried out in that version in two animals (animals R and Z).

In the first version of the task, each trial began with the appearance of a white fixation point at the center of the screen which the animal was required to fixate with its gaze. The task proceeded once the animal maintained fixation continuously for 0.25 s and at least 0.5 s had passed since fixation point onset. If the animal did not fixate within 5 s of fixation point appearance, or broke fixation before 0.5 s had passed since fixation point onset, the trial was counted as an error, moved immediately to the inter-trial interval, and repeated until it was performed successfully. After the fixation requirement was successfully completed, the fixation point disappeared and there was no longer any gaze requirement to advance the task. Simultaneously with fixation point disappearance, the first offer appeared on the screen. After 1 s, the second offer appeared on the screen. After a further 0.5 s, the choice period began. An offer was counted as chosen once the animal held its gaze on it continuously during the choice period for a fixed duration (0.5 s in animals R, B, and P; 0.4 s in animal Z). If the animal did not choose an offer within 5 s of the start of the choice period, the computer randomly selected an offer and the trial proceeded as if the animal had chosen it; this occurred rarely, and these computer-chosen trials as well as error trials were excluded from analysis unless otherwise noted. The size of each offer stimulus was approximately 6 x 5 degrees of visual angle. The locations of the two offers on the screen were randomly selected on each trial from a set of three possible locations (center, 10 degrees left of center, 10 degrees right of center).

Each offer featured a reward distribution that was visually depicted with 4 bars, with the height of each bar proportional to the volume of its juice reward, up to a maximum size Rmax corresponding to the maximum bar height (Table S1). If an offer was chosen, one of its 4 bars was selected uniformly at random as the outcome, and hence the juice volume indicated by that bar was delivered to the animal at the time of outcome delivery. Juice was delivered through a metal spout placed directly in the mouth, to ensure that anticipatory actions such as licking were not required to obtain the reward and did not influence the amount of reward that was received. Each offer also could be informative or non-informative, indicated by its bars having distinct visual textures (we used complex visual textures sourced from images of scenes; for clarity of presentation, these are depicted in the figures as simple colors, i.e. red or blue). The mapping from texture to informativeness was counterbalanced across animals. If an informative offer was chosen, a visual cue was later presented that highlighted the one of its bars that was selected to be the outcome on that trial. If a non-informative offer was chosen, the same visual cue was presented at the same time, but highlighting a random one of the four bars. Thus, choosing an informative offer gave early access to information about the trial’s upcoming reward outcome.

After the choice was made, the unchosen offer disappeared, then after 0.5 s the visual cue appeared on the chosen offer in the form of a black box highlighting one of its bars. After a further 3.5 s a “reveal” occurred in which three of the bars disappeared and only one bar remained, and then after a further 0.5 s the reward was delivered. The remaining bar always indicated the trial’s outcome, and hence, always provided full information about the trial’s upcoming outcome. This was to ensure that both informative and non-informative offers did eventually provide complete information about the trial’s outcome before it was delivered, to ensure that animals always had adequate time to physically prepare to consume the reward. Thus, in this task Tcue, the time of the cue after the choice, was 0.5 s; Treveal, the time of the reveal after the choice, was 4.0 s; and Tadvance, the time the cue appeared in advance of the reveal, was 3.5 s. Finally, the offer disappeared 0.5 s after outcome onset, then a 1.2 s inter-trial interval occurred before the next trial began.

We presented offers with multiple types of reward distributions. For the full details of all reward distributions used in all sessions, see (Table S1). In brief, when randomly generating each offer on each trial, the offer first had its expected reward selected uniformly at random from a pre-specified set of possible expected rewards. To reduce the number of trials with trivial decisions where one offer was much better than the other, for some animals and sessions, the Offer 2’s expected reward was constrained to be within a pre-specified range of Offer 1’s expected reward. Then, the offer had its reward distribution randomly drawn from one of the following types: Safe distributions with 100% probability of delivering a specific reward amount; 25/50/25 distributions with 25, 50, and 25% chances of delivering small, medium, and large reward amounts, respectively, where the medium amount was equal to the mean of the small and large amounts; 50/50 distributions with 50% chances of delivering small or large amounts; and in some animals and sessions, 25/75 distributions with a 25% chance of delivering one reward amount and a 75% chance of delivering a different reward amount. Then, after the offer’s distribution type was selected, its reward range was randomly drawn from a pre-specified set of ranges. For example, in animal Z, uncertain reward distributions could have a range of either 0.35 mL (low risk) or 0.5 mL (high risk); other animals had a wider selection of possible ranges. Finally, the offer’s four possible reward outcomes were each discretized to occur in increments of Rstep between 0 and the pre-specified maximum possible reward size for that animal, Rmax. For example, in animal R, Rstep = 0.02 and Rmax = 0.6, so there were 31 possible reward sizes: 0, 0.02, 0.04, …, 0.6 mL juice, and hence a maximum height bar on the offer stimulus corresponded to a 0.6 mL juice outcome.

In the second version of the task, we augmented the task to manipulate information and reward timing. Specifically, each offer was augmented with a large “clock” in the form of a horizontal bar placed at the bottom of the offer. The horizontal extent of the clock represented the full time duration between choice and the end of the trial (equal to 8 s in this task version). The clock was divided into three sequential segments, representing the segments of time (A) from the choice to the cue, (B) from the cue to the reveal, and (C) from the reveal to the end of the trial. The time of reward delivery was not explicitly indicated but was always exactly 1.1 s after the reveal. The three segments of the clock each had distinct visual textures, which were also distinct for informative vs. non-informative offers (again, for clarity of presentation, these are depicted in figures as simple shades of red and blue). In addition, two thick black vertical lines indicated the time of the cue and the time of the reveal. Finally, to help animals learn how the clock corresponded to real world time, and to help them anticipate the time of each event during each trial, we animated the clock (Fig S2). Specifically, after the choice was made, a “hand of the clock” appeared at the left edge of the clock, in the form of a thick black rectangle whose position on the clock indicated the current time during the trial. The hand moved from left to right at a constant speed as the trial proceeded. When the hand touched the first black vertical line indicating Tcue, an animation played lasting 0.6 s in which the vertical line moved upward to the center of the cued bar and then expanded to become the cue. When the hand touched the second black vertical line indicating Treveal, an animation played lasting 0.6 s in which the vertical line moved upward to the center of the chosen offer and then became an expanding rectangular region with the background color of the screen, ‘erasing’ three of the bars thus leaving only one bar remaining. Finally, when the hand touched the right edge of the clock, the trial ended.

For each offer, the pair of event times, (Tcue, Treveal), was drawn uniformly at random from the set of all possible such pairs meeting the following requirements: Tcue was between 0.4-5.8 s; Treveal was between 1.2-6.7 s; Tcue came at least 0.8 s before Treveal; Tcue and Treveal were both multiples of 0.1 s. In addition, we also altered the statistics of the reward distributions. For the full details of the reward distributions used in all sessions, see (Table S1). In brief, there were two major changes. First, because this version of the task had longer trials, to maintain a high reward rate to keep animals motivated to perform the task, we scaled up the rewards by using a higher maximum reward size Rmax. Second, to make it easier for animals to process and interpret the reward distributions despite the added visual and motivational complexity from the clock, we used a more limited set of reward distributions in some animals.

Finally, during recordings, the multi-attribute information task above was interleaved in alternating blocks with a simpler information anticipation task used in our previous work (*32*). Specifically, after the multi-attribute information task was run for a block of 30correctly performed trials, then the information anticipation task was run for a block of 9 correctly performed trials. The information anticipation task began with the appearance of a large purple fixation point at the center of the screen which the animal was required to fixate with its gaze. Importantly, the fixation point for the anticipation task was clearly visually distinct from the fixation point for the multi-attribute task, so the fixation point indicated to the animal which task was in effect. After the fixation requirement was successfully completed, a single visual fractal CS was presented for 1.5 s, then was replaced by a visual fractal cue for 1.5 s, followed by cue disappearance simultaneous with outcome delivery, followed by a 1.6 s inter-trial interval. The location of the CS on the screen was randomly selected on each trial from the same set of three possible locations used in the multi-attribute task. There were three unique CSs: a Safe CS, a Risky CS, and an Info CS. Importantly, these CSs all yielded exactly the same expected reward volume, but with different levels of risk and access to information. Specifically, each CS was associated with a unique set of two possible cues, one of which was randomly selected to replace that CS on each trial. The Safe CS was replaced by one of two cues which each yielded 100% probability of a medium sized reward. The Risky CS was replaced by one of two cues which each yielded a 50% probability of a large reward (double the size of the medium sized reward) and a 50% probability of no reward. Finally, the Info CS was replaced by either a cue that yielded a 100% probability of a large reward or a cue that yielded a 0% probability of reward. Each block of 9 trials consisted of three presentations of each of the three CSs, in a random order.

### Data acquisition

We recorded neurons in the lateral habenula (LHb), anterior/ventral regions of pallidum (Pal), and subthalamic nucleus (STN). A plastic head holder and plastic recording chamber were fixed to the skull under general anesthesia and sterile surgical conditions. The chambers were tilted laterally by 35-40° and aimed to provide access to the areas of interest. After the animals recovered from surgery they participated in the experiments. Electrode trajectories were determined with a 1 mm-spacing grid system and with the aid of MR images (3T) obtained along the direction of the recording chamber. This MRI-based estimation of neuron recording locations was aided by custom-built software (PyElectrode ^104^). In addition, in order to further verify the location of recording sites, after a subset of experiments the electrode was temporarily fixed in place at the recording site and the electrode tip’s location in the target area was verified by MRI (Fig S7).

Electrophysiological recordings were performed using multi-contact electrodes (Plexon V-probes, 32 channels, 50 μm spacing) inserted through a stainless steel guide tube and advanced by an oil-driven micromanipulator (MO-97A, Narishige). Signal acquisition (including amplification and filtering) was performed using an OmniPlex 40 kHz recording system (Plexon). Spike sorting was performed offline using publicly available software (Kilosort2) to extract clusters from the recordings, followed by manual curation to identify sets of clusters that corresponded to single neurons, the spans of time when they were well isolated, and whether they were located in the regions of interest. Regions of interest were identified based on their electrophysiological characteristics, firing patterns, and locations relative to anatomical landmarks estimated from MRIs and recordings at multiple grid locations. LHb neurons were also identified by having recording depths within ∼0.5 mm of neurons with negatively-signed reward-related activity in response to cues, reveals, and/or reward delivery. A subset of Pal neurons (n=30) were recorded using single-contact glass-coated electrodes (Alpha Omega) or epoxy-coated electrodes (FHC) from which spikes were sorted offline (Plexon Offline Sorter). Neuronal and behavioral analyses were conducted offline in Matlab (Mathworks, Natick, MA).

Eye position was obtained with an infrared video camera (Eyelink, SR Research). Behavioral events and visual stimuli were controlled by Matlab (Mathworks, Natick, MA) with Psychophysics Toolbox extensions. Juice, used as reward, was delivered with a solenoid delivery reward system (CRIST Instruments).

### Electrical stimulation

During electrical stimulation sessions, low-intensity electrical stimulation (50 μA, 400 Hz, 500 ms) was delivered to LHb on a subset of trials. The stimulation strength was chosen based on previous studies in monkeys (*121–124*). Stimulation was delivered in a post-offer time window starting 100 ms after the onset of one of the offers. Specifically, in animal R, stimulation occurred post-Offer 1 on 25% of trials, post-Offer 2 on 25% of trials, and there was no stimulation on the remaining 50% of trials. In animal Z, stimulation occurred post-Offer 1 on 17% of trials, post-Offer 2 on 17% of trials, and there was no offer period stimulation on the remaining 66% of trials. In a subset of trials in Animal Z with no offer period stimulation, stimulation occurred after the choice was made in a pre-cue or pre-reveal time window ending simultaneously with cue or reveal onset, respectively, each occurring on approximately 12.5% of total trials. In animal R, the information anticipation task was not run during stimulation experiments. In animal Z, the information anticipation task was interleaved with the main task and included stimulation on 25% of trials, which was equally likely to occur at one of three times: post-CS starting 100 ms after CS onset, pre-cue ending simultaneously with cue onset, or pre-outcome ending simultaneously with outcome delivery.

### Data analysis

All statistical tests were two-tailed unless otherwise noted. All statistical significance corresponds to p < 0.05 or a 95% confidence interval excluding 0 unless otherwise noted. Neural firing rates in response to each offer were analyzed in a time window 125-500 ms after offer onset. Firing rates for each neuron were converted to normalized firing rates by z-scoring. Specifically, for each neuron, we computed a vector of firing rates representing its full activity throughout the task by binning its data from all times during all trials in non-overlapping 500 ms bins. We then normalized the neuron’s firing rates in our analyses by subtracting the mean of that vector and dividing by the standard deviation of that vector. A neuron was classified as attribute-responsive if the main GLM used to analyze neuronal activity (described below), when fit to its responses to either Offer 1 or Offer 2, yielded a significant effect of at least one attribute (p < 0.05).

#### Psychometric analysis of the value of information

For each individual, we analyzed the subset of trials where one offer was Info and the other was Noinfo, and plotted the choice of Info as a function of the difference in E[r] between the Info and Noinfo offers. We fit the underlying single trial data with a logistic function (specifically a generalized linear model (GLM) for binomial data with a logistic link function). We then estimated the subjective value of information as the function’s indifference point (i.e. the x-coordinate for which the function would produce a 50% choice of info) multiplied by −1, and estimated its standard error by bootstrapping over trials (n=200 bootstraps). To estimate the mean subjective value of information for the human population as a whole, we did the same procedure except fitting the population average of the choice data from the individuals, fitting the logistic function by minimizing the squared error, and bootstrapping over individuals (n=200 bootstraps). To estimate the effect of uncertainty, we performed the same analysis separately for the subsets of trials where both offers were high uncertainty or both offers were low uncertainty (humans: uncertain offers vs certain offers; monkeys: offers with SD[r] greater than or less than the median SD[r] over all offers presented to that animal).

#### Models fit to behavior and neuronal activity

We fit each individual’s binary choice data using GLMs designed to model a standard decision making formulation in which values are computed for each of the two offers, and then the resulting choice probability is a logistic function of the difference in the values:

log(p(choose offer 2) / p(choose offer 1)) = V(offer 2) – V(offer 1)

And the value of each offer *i* is a linear weighted combination of the offer’s vector of n attributes <x_i,1_,x_i,2_,x_i,3_,..,x_i,n_>:

V(offer *i*) = β_1_*x_i,1_ + β_2_*x_i,2_ + … + β_N_*x_i,n_

Thus, the resulting model was a GLM for binomial data with a logistic link function, with the equation: log(p(choose offer 2) / p(choose offer 1)) = β_1_(x_2,1_ - x_1,1_) + β_2_(x_2,2_ - x_1,2_) + … + β_n_(x_2,n_ - x_1,n_)

For analysis of neuronal data, we used an analogous GLM to fit neuronal firing rates in response to the individual offers, for normal data with an identity link function (i.e. equivalent to ordinary linear regression):

Rate(offer *i*) = β_i,0_ + β_i,1_x_i,1_ + β_i,2_x_i,2_ + … + β_i,n_x_i,n_ + ε

…where β_i,0_ is the neuron’s mean or baseline response to the offer and ε is the error term.

For each analysis, we used models tailored to each behavioral task and specific analysis question at hand, producing a total of 26 models as described below (see Tables S2 and S3 for full details and comparisons of all parameters used by all models). For example, attributes related to time were only included in models for tasks that manipulated time-related variables, and the significance of these time effects on the value of information was evaluated by performing model comparisons between alternate models where Info x Time interaction terms were included vs excluded. There were also several aspects of the model fitting procedure that held across models. First, to permit more interpretable comparisons between attributes, unless otherwise noted, all attributes that were not binary were standardized by z-scoring. Second, in addition to the terms described below related to the values of the offers, all models also included terms for response biases, including the location in space of each offer and the sequential order in which the offers were presented. For monkey behavioral data, this included binary attributes representing whether or not the offer was on the left side of the screen, whether or not the offer was on the right side of the screen, whether the offer was Offer 1 or Offer 2. For monkey neuronal data, this included binary attributes representing whether or not the offer was on the left side of the screen, whether or not the offer was on the right side of the screen, and a constant term representing the neuron’s average level of activity; no term for sequential order was included because the model was fit separately for neural responses to Offer 1 and Offer 2. For human data, this consisted of one binary attribute representing whether the offer was on the right side of the screen. Third, in certain cases we used a fitted behavioral model to derive an estimate of the subjective values that an individual assigned to the offers. To do this, we simply plugged the fitted weights (β_i,1_, …, β_i,n_) and that offer’s attributes (x_i,1_, …, x_i,n_) into the equation for V(offer *i*) above. This produced an estimate of the subjective value that the individual assigned to each offer presented on each trial, in units of the offer’s log odds of influence over the choice.

To compare a set of models and judge which of them fit best, we compared the models in terms of their log likelihood. For human population data, the log likelihood was summed over individuals; for neuronal population data, it was summed over neurons. We used a shuffling correction to control for the possibility that certain models could fit the data with higher likelihood simply due to having more parameters (or due to any other aspect of the structure of the models that could give some of them more power to explain data by chance). Specifically, we fit each GLM on 10 shuffled versions of the dataset in which the variable to be predicted (behavioral models: choice of Offer 1 vs Offer 2; neural models: firing rate in response to the offer) was shuffled randomly across trials, and then defined the shuffle-corrected log likelihood as the log likelihood from the fit to the real data minus the mean log likelihood of the fits to the shuffled datasets. Finally, to test whether any given model had higher shuffle-corrected log likelihood than any other given model, we repeated the above procedure 2000times on 2000 bootstrap datasets generated by randomly resampling the original dataset’s individuals (for human population data), trials (for single monkey data), or neurons (for monkey neuronal data) with replacement. We used this to calculate the bootstrap standard error of each model’s shuffle-corrected log likelihood, and the 95% bootstrap confidence interval of the difference between the shuffle-corrected log likelihoods for each pair of models. One model was deemed better than another model if the confidence interval excluded 0. Model interpretations were also aided by tests of whether key parameters were significant or were significantly different from each other (using the linhyptest function in matlab).

The first set of analyses focused on behavior during the first version of multi-attribute information task for humans and monkeys. The main model for this analysis had attributes for expected reward (E[r]), reward uncertainty (Unc[r]), Informativeness (Info), and the interactions between Info and the other variables (Info x E[r] and Info x Unc[r]). In this set of analyses the Info attribute was centered at 0 by setting it to +0.5 for Info offers and −0.5 for Noinfo offers. We fit separate versions of the model using different hypothesized forms of uncertainty: standard deviation, range, and entropy (*51*), as follows:

SD[r] = (∑ p(r) (r – E[r])^2^)^0.5^

Range[r] = max(r) – min(r)

Entropy[r] = - ∑ p(r) log_2_(p(r))

As a control, we also fit analogous models without any uncertainty-related attributes (i.e. Unc[r] and Info x Unc[r]). We then used the model comparison procedure described above to compare these models. The procedure consistently selected SD[r] as the best measure for both humans and monkeys, so we used SD[r] as the default uncertainty measure for all further models unless otherwise noted.

The second set of analyses was a more detailed characterization of the form of uncertainty that motivates information seeking. We fit models that replaced each uncertainty-related attribute (i.e. Unc[r] and Info x Unc[r]) with a set of separate attributes corresponding to the separate types of reward distributions used in the task. For example, in datasets with two reward distribution types, 25/50/25 and 50/50, the single Unc[r] attribute was replaced by two binary attributes indicating whether the offer was 25/50/25 and whether it was 50/50; similarly, the Info x Unc[r] attribute was replaced by two attributes that were the former two attributes multiplied by Info. We also fit models that further subdivided each reward distribution type into offers of that type that were relatively high vs. low uncertainty (Fig S4, “Distrib UncType Detailed” model in Table S3).

The remaining analyses focused on tasks that manipulated event timing (the second version of the tasks) and on relating neuronal activity to behavior. For the human timing task, the behavioral model had two attributes, E[r] and EarlyInfo, where EarlyInfo indicated whether the offer yielded early information vs. late information. For the monkey timing task there were a much larger number of possible variables to consider, since it included manipulations of both Tcue and Trew in addition to all of the other task variables described above.

Therefore, we used a model selection procedure to identify the core set of attributes that most strongly governed the subjective value of offers across monkeys. The goal was to identify attributes that explained a large fraction of the variance in behavior in the very large behavioral datasets collected for each monkey, while still having a limited number of attributes so that the same model could be fit practically to the smaller datasets collected when recording each individual neuron. Importantly, these attributes were selected purely based on fits to behavioral data, independent of neuronal data. This selection procedure was performed on a preliminary behavioral dataset that included all three animals that provided data for the second version of the monkey task. In the procedure, we first started with a large set of possible attributes to include in the model (n=18), consisting of: E[r], SD[r], and Info; all pairwise interactions between those three variables; Tout and its pairwise interactions with those three variables; Tadvance and its pairwise interactions with those three variables; and all three-way interactions between Info, either Tout or Tadvance, and either E[r] or SD[r]. Then for each animal, we ranked the attributes by their importance in explaining behavior using an iterative procedure. We started by fitting the behavioral data with an empty model without any of those attributes (i.e. only including the standard bias terms), and measured its cross-validated log likelihood (10-fold cross-validation). We then iteratively added individual attributes to the model, at each step adding the single attribute that improved the cross-validated log likelihood the most, and stopping when no attributes produced improvements. Thus, for each animal, this produced a sequence of attributes ranked by their marginal importance in explaining choice behavior. This also placed lower and upper bounds on the ability of these models to explain the data, defined by the cross-validated log likelihood of the initial empty model vs. the final model. Finally, based on these results, we selected the subset of attributes that was sufficient to raise the cross-validated log likelihood from the lower bound up to 99.9% of the way to the upper bound, across all animals. This produced the following 10 core attributes: E[r], Tout, SD[r], E[r] x Tout, E[r] x SD[r], Info, Info x Tout, Info x SD[r], Info x Tadvance, Info x Tadvance x SD[r]. We then defined a model with these attributes as the main model for the second version of the monkey task, and used it as the basis for our analysis of both monkey behavior and neuronal activity in this task.

Specifically, just as we had done for the behavioral models of the first version of the task, we used the main model with the above attributes to fit data from the second version of the task. As before, we compared the model fit between the main model described above, which uses SD[r], versus alternate models using either Range[r], Entropy[r], or no uncertainty-related terms. Similarly, as before, we compared the fitted weights of the Info x SD[r] and Info x E[r] attributes to judge their relative effects; note that for this analysis we used an alternate version of the model that included Info x E[r] as an additional term, since the Info x E[r] interaction term was not identified as a core attribute by the procedure above (consistent with its relatively small effect on behavior in this task; Figure 1). Next, to investigate how information value is computed using uncertainty and time, we did a model comparison between the full model, a model that removed attributes with interactions between Info and Unc[r], a model that removed attributes with interactions between Info and time-related variables (Tout and Tadvance), and a model that removed attributes with both types of interactions.

Next, for a more detailed characterization of how information value depends on the timing of cues and outcomes, we fit a version of the model where all terms with interactions between Info and time-related variables were removed, and instead, three attributes were included representing the interaction between info and the presence of specific timing properties in the offer. Specifically, we defined cues as ‘early’ if Tcue was within the 0^th^-33^rd^ percentiles of possible cue times, and ‘late’ if Tcue was within the 66^th^-100^th^ percentiles of possible cue times. We also defined outcomes as ‘early’ and ‘late’ with an analogous rule. We then included three attributes to the model: Info x (early cue, early outcome), Info x (early cue, late outcome), and Info x (late cue, late outcome); where (x,y) was 1 for offers that met both the specified cue and outcome timing conditions, and 0 otherwise.

Finally, to compare across species, we computed the effect of Early vs Late Info on subjective value in animals and humans, in units of log odds of choice. To quantify this for each human, we used the fitted GLM weight of EarlyInfo. To quantify this for each monkey, we used the main model’s GLM fit to derive the subjective value of each offer, and computed the mean difference in offer subjective value that resulted from info cues being presented early vs. late. To produce a measure a closely analogous as possible to the human data, this analysis only used the subset of offers that were most closely analogous to those used in human task (i.e. offers with high reward uncertainty and late outcome delivery, as defined above). We then defined the effect of interest as the difference between the mean derived value of Info offers vs. Noinfo offers with early cue delivery, minus the difference between the mean derived value of Info vs Noinfo offers with late cue delivery. Error bars and confidence intervals were computed with bootstrapping (n=2000 bootstraps).

#### Value coding index

We defined a value coding index to quantify the extent to which the attribute-related offer responses of each individual neuron aligned with our behaviorally-derived estimate of how animals weighted those attributes to compute the subjective value of those offers. To do this, we first fit each neuron using the main model described above, which we will refer to here as the “attribute model” because it allows the neuron to have separate weights for each of the 10 separate attributes described above that influenced the animal’s choices. We then fit each neuron with a new model called the “value model”, which only had a single attribute, “Value”, equal to the estimated subjective value of the offer derived from the main model (using the procedure described above). Thus, the value model represents the hypothesis that the neuron’s attribute-related activity is fully explained by encoding of a single variable, the offer’s fully integrated subjective value; or equivalently, that the neuron’s weighting of the offer’s many attributes to govern its response, is perfectly correlated with the animal’s weighting of the offer’s many attributes to compute subjective value and guide its decisions.

Finally, we quantified what fraction of the above-chance variance in the neuron’s activity that was explained by the attribute model could also be explained by the value model. To do this, we used a measure based on R-squared (R^2^) to quantify the fraction of the response variance that each model explained. Specifically, to correct for the fact that the attribute model had more parameters than the value model, we computed the shuffle-corrected R-squared (R^2^corr), defined as the R^2^ computed on the neuron’s real data minus the mean of the R^2^ computed on 100 shuffled datasets in which the neural responses to the offers were shuffled across trials. Thus, R^2^corr quantified the fraction of variance of neural responses that the model explained, above and beyond the chance level expected under the null hypothesis that attributes and values had no influence on the neuron’s activity. We then calculated the mean R^2^corr for each model as the average of the R^2^corr that were computed from separate fits of that model to the neuron’s Offer 1 and Offer 2 responses. Finally, we defined the value coding index with the equation:

ValueCodingIndex = *c*((mean R^2^corr for value model) / (mean R^2^corr for attribute model))

…where the function *c*(*x*) = max(0,min(1,*x*)) clamps the index between 0 and 1 (for rare cases where the index was slightly < 0 or > 1 such as due to small variations in the shuffling used for the shuffle correction; n=8 LHb, n=8 Pal, n=5 STN). Importantly, because the value coding index is expressed as a ratio relative to the corrected variance explained by the attribute model, it is only meaningful and reliably estimated for neurons for which a meaningful fraction of the variance was explained by the attribute model. Therefore, we only computed this index for neurons that had strong attribute effects, which we defined as neurons for which the mean R^2^corr for attribute model was greater than or equal to 0.1 and that were significantly attribute responsive.

#### Analysis of choice-predictive activity

We computed a choice predictive index as a measure of how variations in LHb neuronal activity above and beyond those accounted for by our neuronal model, were predictive of variations in choice behavior above and beyond those accounted for by our behavioral model. To do this, we first fit the behavioral choice data with the main model. To ensure that the behavioral fit reflected the animal’s behavioral policy as closely as possible during recording of the neuron, we used a fit that was restricted to the subset of the behavioral data that was collected during the recording of that neuron (we obtained similar results when using the behavioral fits to the full behavioral dataset). Next, we fit the neuronal offer responses with a version of the main model. We did separate fits for the neuronal responses to Offer 1 and to Offer 2. To allow comparison across neurons, rather than fitting raw firing rates, we fit “normalized value signals”, defined as the normalized firing rate that has been sign-flipped based on the that neuron’s fitted sign of value coding (from the value coding index analysis described above). Thus, on average, more positive normalized value signals correspond to activity reflecting higher offer values. In addition, to control for the possibility that variations in the neuronal response to the currently fitted offer might be influenced by the properties of the other offer, this was done based on a fit of a versions of the model that, in addition to the terms representing how the response to the currently fitted offer response depended on each attribute of that offer, also included an analogous set of terms for how the response to the currently fitted offer depended on each attribute of the other offer. Thus, for each trial, these fits provided us with a predicted choice probability and predicted normalized firing rates for Offer 1 and Offer 2. Using these, for each offer *i*, we computed the behavioral residual Δbehavioral*_i_* and the neural residual Δneural*_i_*, using the following equations:

Δbehavioral*_i_* = (Chose Offer *i*?) – (predicted p(choose Offer *i*))

Δneural*_i_* = (normalized value signal for Offer *i*) – (predicted normalized value signal for Offer *i*)

Finally, to improve the power of our analysis to relate variations in neurons to variations in behavior, we excluded the subset of trials with offer pairs that produced highly deterministic choice behavior (e.g. due to one offer having much higher value than the other), and hence where even very large variations in neuronal activity in one direction were only mathematically capable of being associated with very small behavioral residuals in that direction. Specifically, we excluded trials in which the behavioral model predicted that the animal would choose a specific offer with > 95% probability and that prediction was correct (i.e. trials in which Δbehavioral*_i_* was bounded to be <= +0.05 or <= −0.05). We then defined the choice predictive index for each Offer *i* as the Spearman’s rank correlation between the behavioral and neural residuals:

ChoicePredictiveIndex*_i_* = corr(Δbehavioral*_i_*, Δneural*_i_*)

…where the behavioral residual Δbehavioral*_i_* = (Chose Offer *i*?) – (predicted p(choose Offer *i*)), and the neural residual Δneural*_i_* = (normalized value signal for Offer *i*) – (predicted normalized value signal for Offer *i*). In addition, to pool neural responses over the two offers to establish a total prediction about each trial’s choice, we computed a total choice predictive index for each neuron as the correlation between the residual choice of Offer 2 and the difference in neural residuals reflecting the net difference in normalized residual value signal favoring Offer 2:

ChoicePredictiveIndex = corr(Δbehavioral*_2_*, Δneural*_2_* – Δneural*_1_*).

To examine the relation between behavioral and neural residuals at the population level, we calculated a population average relationship between residual value signals and residual choice, separately for Offer 1 and Offer 2. Specifically, for each neuron, we normalized its residual value signals by dividing the residual value signal on each trial by the SD of its residual value signals over all trials, and then clamping them between −4 and +4. Then we plotted the relationship between, and calculated the correlation between, the normalized residual value signals and residual choices, pooling over all neurons and all trials. To test for a difference between Offer 1 and Offer 2, we performed the above procedure on bootstrapped datasets (n=200 bootstraps over neurons) and tested whether the bootstrap confidence interval of the difference between the correlations for Offer 1 and Offer 2 excluded 0.

As a further test of whether LHb residual value signals could be used to predict choice behavior, we asked whether they could be used to improve the performance of our behavioral models. First, for each neuron, we used the fit of the main behavioral model described above to derive an estimate of the subjective values of Offer 1 and Offer 2 on each trial (“V1” and “V2”, respectively), as described above. Then, we fit a new behavioral model (Table S3) that had two attributes: (1) “V2-V1”, equal to the difference in subjective value between the two offers, and (2) “LHbResid2-LHbResid1”, equal to the difference in normalized residual value signals (Δneural*_2_* – Δneural*_1_*). To further separate the influences of Offer 1 vs Offer 2, we also fit alternate models in which LHbResid1 and LHbResid2 were included as separate attributes, to allow them to be fitted with separate weights for predicting behavioral choices (Table S3).

#### Analysis of reward prediction error coding

To quantify each neuron’s coding of conventional juice reward prediction errors, we computed a reward prediction error index (RPEindex) using the following approach. First, we defined the RPE on each correctly performed trial as the difference between the delivered reward and the chosen offer’s E[r]. Next, we selected the subsets of trials which fell into each of four categories, based on the 2×2 combinations of the chosen offer’s informativeness (Info or Noinfo) and the resulting RPE (Pos or Neg, corresponding to RPEs > +0.1 mL juice or < −0.1 mL juice). We then analyzed single trial firing rates on these trials after two task events where RPEs commonly occurred in our task: a post-cue time window 150-750 ms after cue onset, and a post-reveal time window 150-750 ms after reveal onset. We then used this activity to compute separate measures of the strength of RPE-related responses in each of those time windows (RPEcue and RPErev), described below. Importantly, RPEcue is focused on Info trials while RPErev is focused on Noinfo trials. This is because the cue only provides new information about the outcome on Info trials, and hence should only produce RPEs on Info trials; on Noinfo trials, the cue is non-informative and should not produce RPEs (*18, 35*). Similarly, the reveal only provides new information about the outcome on Noinfo trials, and hence should only produce RPEs on Noinfo trials; on Info trials, the cue already indicated the outcome in advance, so the reveal provides no new information and should not produce RPEs (*18, 35*). Indeed, those were exactly the times in the task when our LHb neurons produced prominent RPE signals. Thus, RPEcue was designed to reflect the degree to which a neuron’s cue-period activity reflects RPEs on Info trials (when RPEs should occur during the cue period) vs. Noinfo trials (when they should not occur during the cue period), while RPErev for the reveal period did so in the analogous manner for Noinfo trials vs. Info trials, as follows:

RPEcue = ROC(InfoPos post-cue, InfoNeg post-cue) - ROC(NoinfoPos post-cue, NoinfoNeg post-cue) RPErev = ROC(NoinfoPos post-rev, NoinfoNeg post-rev) - ROC(InfoPos post-rev, InfoNeg post-rev)

…where ROC(x,y) is the ROC area (area under the receiver operating characteristic (*125*)) between the set of single trial firing rates in condition x vs. condition y, such that the ROC area is > 0.5 if condition x generally has higher activity than condition y, and < 0.5 if condition x generally has less activity than condition y, and 0.5 if the distributions are the same. Significance was assessed by permutation tests (p < 0.05, n=400 permutations) in which each of these measures was compared with the distribution of the same measure computed on permuted datasets in which firing rates were shuffled between InfoPos and NoinfoPos and between InfoNeg and NoinfoNeg, reflecting the null hypothesis that activity was not different between the conditions during which RPEs should occur vs. should not occur. Finally, the total RPEIndex was computed as the mean of these two measures:

RPEIndex = (RPEcue + RPErev) / 2

…and its p-value was computed from the p-values of the two constituent measures using Fisher’s combination test (*126, 127*).

#### Analysis of overlap between coding properties

For each neural population we tested the overlap between the three coding properties measured by the indexes described above: ValueCodingIndex, ChoicePredictiveIndex, and RPEIndex. We classified cells as strongly value coding if they had ValueCodingIndex > 0.6, and as significantly choice predictive or RPE coding if the respective indexes were significant (p < 0.05). This analysis included all attribute-responsive neurons for which all three coding properties could be computed. We classified neurons as “fully combined” if they passed all three criteria and all three had identical coding signs. For example, if a neuron had all three coding properties with negative signs, it would have offer responses negatively related to offer subjective value, lower firing rate during offer responses predicting greater choice of that offer, and lower firing rate during cue/reveal responses associated with more positive RPEs. We used permutation tests (n=200,000 permutations in which each index was independently shuffled across neurons) to test whether the proportion of “fully combined” neurons in an area was greater than chance, and whether the difference of that proportion between areas was significantly greater than chance. Finally, we plotted a venn diagram depicting these neurons, in which the area of each of the three main circles was proportional to the percentage of these neurons with the corresponding coding property, and the intersections between pairs or triples of circles represented neurons that had pairs or triples of coding properties and coded them with identical signs. The diagram was optimized by fixing the areas of each circle and then adjusting the centers of the circles to minimize the total error between the desired and displayed areas of the regions representing the intersections between sets of circles (with regions that did not contain any neurons treated as having a minimal area corresponding to 0.1% of these neurons).

#### Analysis of peri-stimulus activity

To plot the cross-validated timecourse of normalized activity related to the interaction between Info and Uncertainty, we used the following procedure. First, separately for each attribute-responsive neuron and separately for Offer 1 and Offer 2, we classified all offers into the 2×2 categories of (Info, Noinfo) x (Uncertain, Safe). Then we split this data into two cross-validation folds, 1 and 2, which included odd-numbered and even-numbered trials from within each these categories, respectively. For each fold *i*, we computed its Info x Uncertainty effect, InfoXUnc_i_, as:

InfoXUnc*_i_* = (Rate*_i_*(Info, Uncertain) – Rate*_i_*(Noinfo, Uncertain)) - (Rate*_i_*(Info, Safe) – Rate*_i_*(Noinfo, Safe))

…where Rate*_i_*(*offer type*) is the neuron’s mean normalized firing rate in response to the offers of the specified type using the subset of trials that were included in fold *i* and the standard analysis window for responses to that offer. Finally, we computed the total cross-validated effect InfoXUnc, by taking the mean of the effects in the two folds, after multiplying each fold’s effect by the ‘uncertainty coding sign’ derived from the other fold’s effect:

InfoXUnc = (sign(InfoXUnc*_1_*)*InfoXUnc*_2_* + sign(InfoXUnc*_2_*)*InfoXUnc*_1_*) / 2

This provided a cross-validated measure of the total strength of the Info x Uncertainty interaction in units of normalized activity, such that a more positive effect indicates a stronger interaction, regardless whether the underlying effect on firing rate had a positive sign (more positive info effect under uncertainty) or a negative sign (more negative info effect under uncertainty). Specifically, if the neuron had a true effect that could be measured with a consistent sign in both folds, then InfoXUnc would be positive (regardless of whether the sign of the underlying effect reflected a positive or negative influence on firing rate). Whereas, under the null hypothesis that the neuron had no true effect, then InfoXUnc would have an expected value of 0, because each fold would be equally likely to be multiplied by either a negative or positive sign. For each neuron, we computed this InfoXUnc separately for Offer 1 and Offer 2 and then averaged the two to get the overall InfoXUnc effect for that neuron. We then tested whether the median InfoXUnc effect across the population of neurons was different from 0 (signed-rank test). The above analysis was restricted to the mean normalized firing rates in the standard offer response window. We also plotted the full timecourse of the InfoXUnc effect by computing it using the full spike density functions from each cross-validation fold and each one of the 2×2 conditions (smoothed with a Gaussian kernel, σ = 40 ms), and while holding sign(InfoXUnc_1_) and sign(InfoXUnc*_2_*) fixed at the values used in the analysis above (so that the full timecourse of each neuron’s activity in each fold was normalized using a consistent sign).

To plot the cross-validated timecourse of normalized activity related to the interaction between Info and specific types of uncertainty (InfoXUncType), we used the same procedure with two modifications. First, instead of comparing Uncertain vs Certain offers, we compared Uncertain 50/50 vs Uncertain 25/50/25 offers. Second, we used the same fixed effect signs that had been computed for the previous analysis (sign(InfoXUnc*_i_*)). This ensured that the results were directly comparable to the previous analysis, and also served as an additional cross-validation step, by ensuring that the ‘uncertainty coding sign’ used for each neuron and fold in this analysis was selected based on the previous independent analysis that had no knowledge of the 50/50 vs 25/50/25 comparison. Thus the equations were:

InfoXUncType*_i_* = (Rate*_i_*(Info, 50/50) – Rate*_i_*(Noinfo, 50/50)) - (Rate*_i_*(Info, 25/50/25) – Rate*_i_*(Noinfo, 25/50/25))

InfoXUncType = (sign(InfoXUnc*_1_*)*InfoXUncType*_2_* + sign(InfoXUnc*_2_*)*InfoXUncType*_1_*) / 2

To plot the cross-validated timecourse of normalized activity related to the interactions between Info and time-related variables we used a slightly different procedure, to accommodate the fact that there were three different Info x Time attributes to measure (Info x Tout, Info x Tadvance, Info x Tadvance x Unc[r]), and that individual neurons might only have activity related to a subset of these effects (e.g. in Pal). Specifically, separately for each neuron and for Offer 1 and Offer 2, we fit the main model for the second version of the monkey task to the normalized activity in response to that offer, using the standard analysis time window for offer responses. We then used these fits to calculate a cross-validated measure of the summed strength of the fitted weights for subset of the Info x Time attributes that the neuron encoded, by performing cross-validation across the two offers to select which subset of attributes the neuron encoded and to select their coding signs. Specifically, we will denote the three attributes as attributes 1, 2, and 3; their fitted weights in response to each offer *o* as *w_o1_*, *w_o2_*, *w_o3_*; and their p-values as *p_o1_*, *p_o2_*, *p_o3_*. We then defined the InfoXTime effect for attribute *a* as the attribute’s mean fitted weight during offers for which that attribute was encoded. Crucially, this effect was cross-validated because both the attribute’s coding sign, and whether it was significantly encoded during each offer, were selected solely on the basis of its responses to the other offer, as follows:

InfoXTime*_a_* = (*w_1a_*\*sign(*w_2a_*)*sig(*p_2a_*) + *w_2a_*\*sign(*w_1a_*)*sig(*p_1a_*)) / max(1,sig(*p_1a_*) + sig(*p_2a_*))

…where the significance indicator function sig(*p*) is 1 if *p* < 0.05 and 0 otherwise. As in the previous analyses, if there is a true effect that is consistently measurable as significant with a consistent sign then InfoXTime*_a_* would be positive, while under the null hypothesis of no true coding InfoXTime*_a_* would have an expected value of 0. Finally, we computed the total InfoXTime effect by summing the InfoXTime effects for the three attributes:

InfoXTime = ∑_a_ InfoXTime*_a_*

As in the previous analysis, we then tested whether the median InfoXTime effect across the population of neurons was different from 0 (signed-rank test). The above analysis was restricted to the standard offer response window. We also plotted the full timecourse of the InfoXTime effect using a sliding analysis in which separate fits were done in 4 ms increments on the set of single trial spike density functions (converted to normalized firing rate and smoothed with a Gaussian kernel, σ = 40 ms).

#### Analysis of electrical stimulation

As a first pass, we used psychometrics to test whether LHb electrical stimulation after the onset of an offer influenced the subsequent choice of that offer. We calculated psychometric curves representing the percent choice of Offer 2 as a function of the estimated difference in subjective value between Offer 2 and Offer 1. To do this, for each animal we first fit the main model of the second version of the monkey task to the subset of choice data collected from stimulation sessions (Table S1), using only trials in which there was no stimulation during the offers. Based on this model fit, we used the procedures described above to derive an estimate of the subjective value of Offer 1 and Offer 2 (V1 and V2), and the difference between the two offer values (V2-V1), on each trial. All of these estimated values were in units of log odds of influencing choice. Importantly, we derived these value estimates even for trials in which stimulation was applied; thus, this gave us an estimate of what the difference in estimated subjective value of the two offers “should have been” on those trials if LHb stimulation had not been applied. We then plotted the percent choice of Offer 2 as a function of (V2-V1) separately for trials with no stimulation, trials with Offer 1 stimulation, and trials with Offer 2 stimulation. For each curve, we estimated the indifference point using the same approach described above (i.e. by fitting the underlying choice data with a logistic function and using the confint function in Matlab to compute its confidence interval). Finally, we tested whether the 95% confidence interval of the indifference point excluded 0.

Next, we quantified the stimulation effect more precisely by using a modeling approach. First, to measure the mean effect of stimulation regardless of whether it occurred during Offer 1 or Offer 2, we fit each animal’s total behavioral dataset during stimulation sessions with a model with three attributes: (V2-V1), Stim1, and Stim2 (Table S3). The effect size and significance of stimulation was defined as the fitted weight and significance of the (Stim2-Stim1) attribute. In addition, to measure the stimulation effect separately for Offer 1 and Offer 2, we fit a version of the model with separate weights for Stim1 and Stim2. In addition, to measure the LHb stimulation effect from each individual session, we fit this model to each individual session separately.

Finally, we used a model comparison approach to ask whether the LHb stimulation effect during Offer 2 was well fit as a simple main effect of stimulation (consistent with stimulation subtracting subjective value from the stimulated offer). Alternately, stimulation might have no effect; might interact with the value of the offer (e.g. larger effect for offers with high value); might interact with the difference between the values of the offers (e.g. larger effect when Offer 2 was higher value than Offer 1); or might interact with specific attributes of the offers (e.g. larger effect for Info offers). To test this for each animal, we fit the full dataset including all stimulation sessions with a set of models representing each of those possibilities (Table S3,S4). All models had the attribute (V2-V1), representing the difference in estimated value between the offers and computed in the manner listed above. The model “StimPred None” had no other attributes; the model “StimPred Stim2” and all subsequent models listed here had an attribute for Stim2; the model “StimPred Stim2xV2-V1” also had an additional attribute for the interaction between Stim2 and the value difference V2-V1; and “StimPred Stim2xV2” instead had an additional attribute for the interaction between Stim2 and V2. Finally, the model “StimPred Stim2xAttrib” was formulated using an interative model selection procedure to select the subset of possible interactions between Stim2 and specific attributes of Offer 2 that significantly improved the model’s fit. Specifically, we started with the model “StimPred Stim2” as the base model for the procedure. We then modified versions of the base model, each of which was the same but with one additional attribute, representing the interaction between Stim2 and the corresponding one of the 10original attributes from the main model for the second version of the task. If none of these modified models had a significant effect of the added attribute, we terminated the procedure. If at least one of the modified models had a significant effect of the added attribute, we selected the modified model whose added attribute had the lowest p-value, designated it as a the new base model, and repeated this procedure. We continued this procedure until it terminated, thus producing a model in which each added Stim2 x attribute interaction had significantly improved the fit. We evaluated all of these models using the shuffle-corrected log likelihood described above, to control for the fact that they had different power to fit the data even under the null hypothesis that there were no true effects (either due to having different numbers of parameters, or due to parameters being selected to significantly improve the fit (for the final “StimPred Stim2xAttrib” model)). Finally, to test the reliability of the difference between the models in shuffle-corrected log likelihood, we used a bootstrapping procedure: we applied the entire procedure described above to 10000 separate bootstrap datasets in which trials were resampled with replacement, then tested if the 95% bootstrap confidence interval of the difference between shuffle-corrected log likelihoods excluded 0.

## Supplemental Material

The supplemental material includes Figures S1-S7, Tables S1-S4 and Supplemental Discussion.

**Figure S1.**
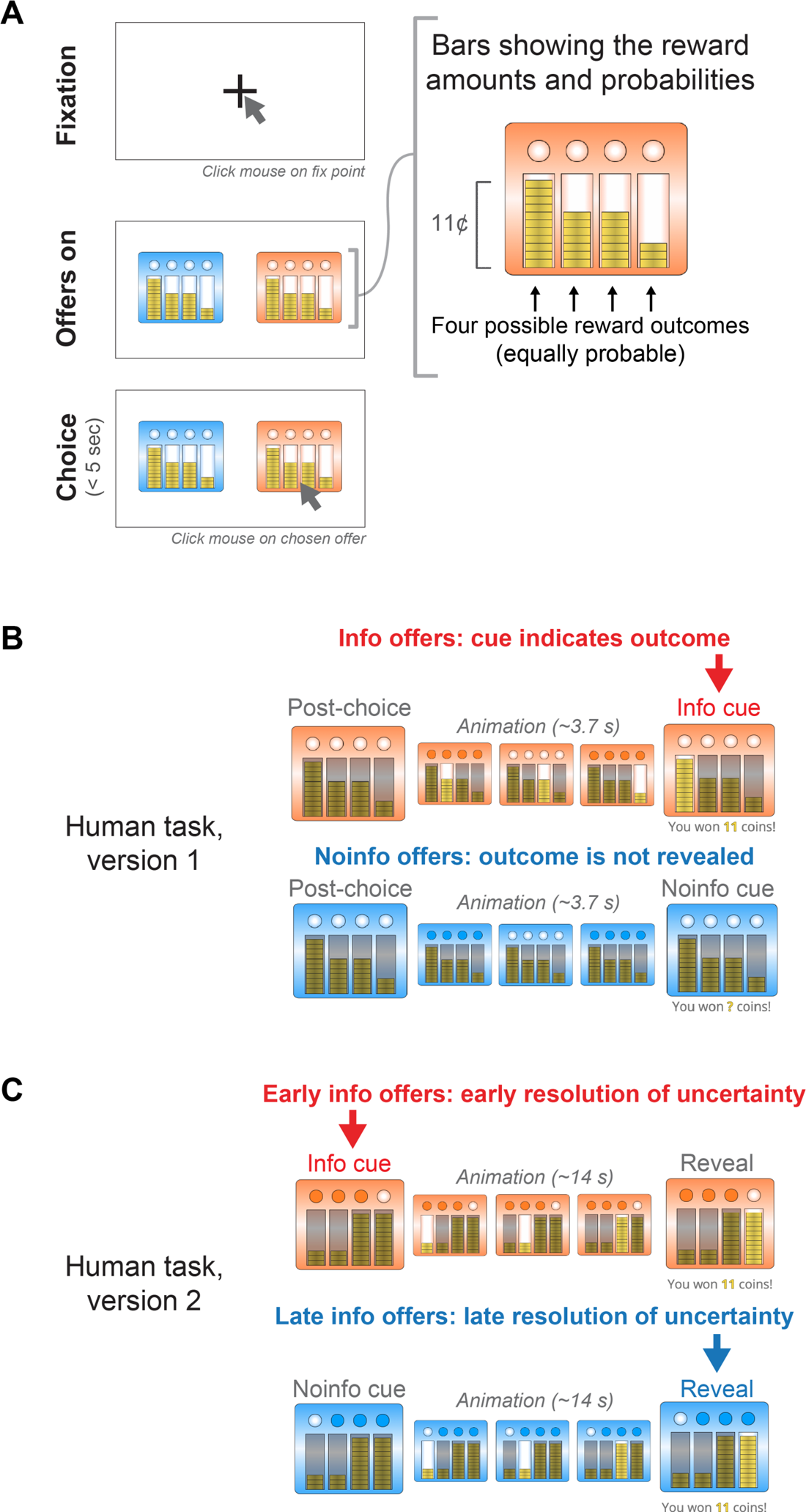
The multi-attribute information choice tasks for humans. (A) After clicking on a fixation point, participants were presented with two offers and chose between them by clicking on the chosen offer. Same as Figure 1A. (B) In the first version of the task, after choosing an Info offer (top), an ∼3.7 s long animation would play in which a single coin stack at a time would be ‘lit up’ for a period of 0.19 s, with the ‘lit’ location initially being a random stack and then cycling sequentially among the stacks from left to right (wrapping around from the rightmost stack to the leftmost stack). The animation ended after 4 full cycles had been completed and the currently lit bar was the one the computer selected to be delivered, at which point text appeared indicating the number of coins that had been won (e.g. “You won 11 coins!”). In addition, during the animation the small circular lights at the top of the offer all alternated between ‘lit’ or ‘not lit’ each time the lit stack changed. After choosing a Noinfo offer (bottom), a matched ∼3.7 s long animation would play, in which the small circular lights at the top of the offer all alternated between ‘lit’ and ‘dark’ just as they did for Info offers, but in which no coin stacks were lit, and the text at the end did not reveal the number of coins that had been won (e.g. “You woncoins!”). (C) In the second version of the task, after choosing an Early Info offer (top), an animation was played which was very similar to the Info animation in the first version of the task, except that it ended after 8 cycles with each animation frame lasting 0.38 s (for a total duration of ∼14 s from choice to final reveal), and immediately after the choice and 1 s before the cycling animation began, one of the four small circular lights at the top of the offer became ‘lit’ and the other three lights became ‘dark’, which accurately indicated the stack that the computer had selected to be delivered on that trial (“Info cue”). Thus if participants chose Early Info, they got early information to resolve their uncertainty about which of the four possible outcomes they would receive. After choosing a Late Info offer (bottom), the same animation occurred except that the ‘lit’ circular light at the top of the offer was randomized and hence did not provide any information about the outcome (“Noinfo cue”). Thus participants only got information to resolve their uncertainty at a later time, from the final ‘lit’ coin stack and the text message indicating their winnings.

**Figure S2.**
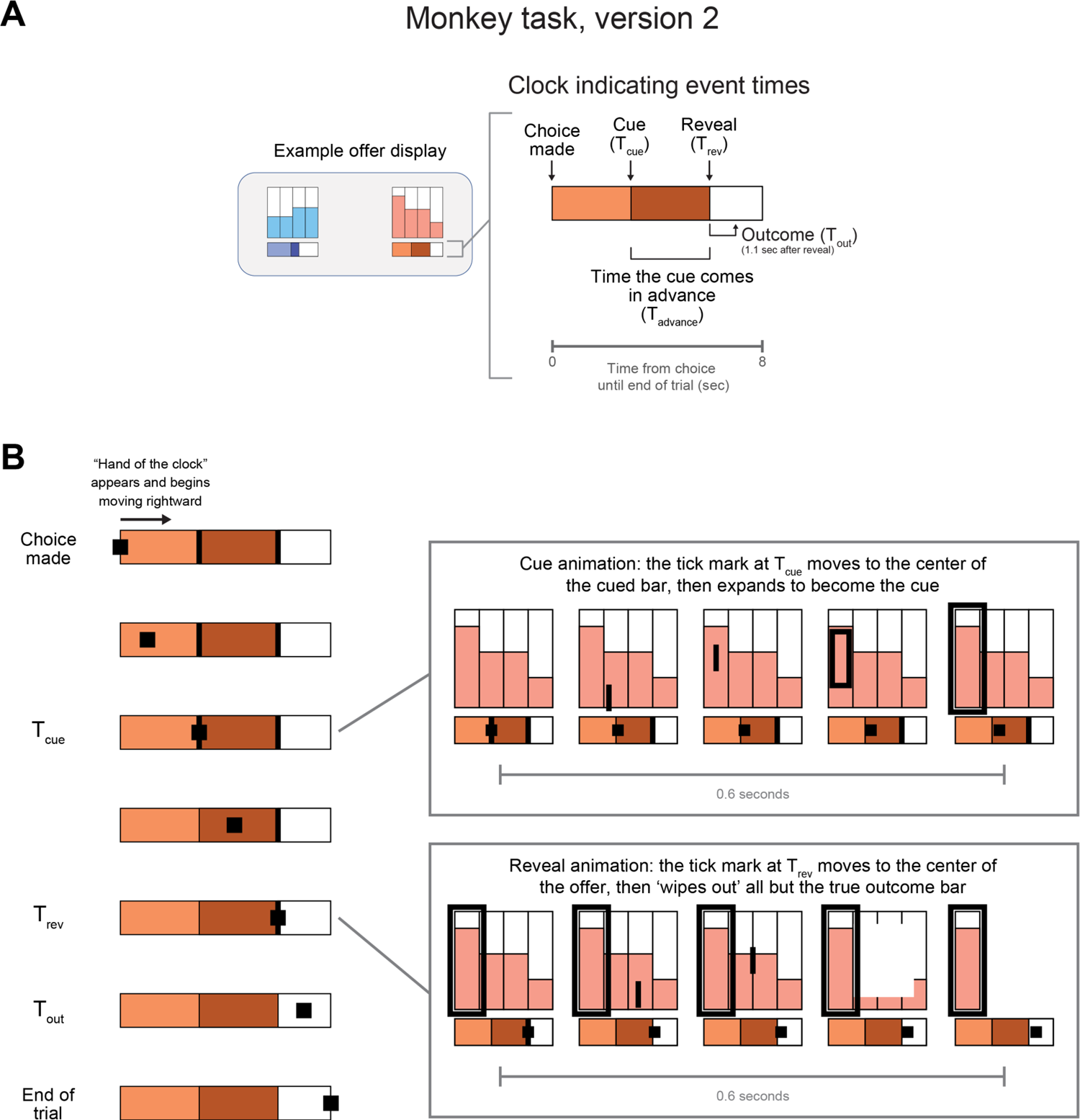
The second version of the multi-attribute information choice task for monkeys, manipulating cue and reward timing. (A) Example Info and Noinfo offer stimuli. Note that the stimuli shown on the screen were complex visual textures sourced from scenes (Methods), which here are represented as shades of red and blue. The top part of the offer indicates the reward distribution (same as the first version of the task). The bottom part is a horizontal bar indicating the timing of task events if the offer is chosen. The leftmost edge of the bar indicates the time of the choice, and the rightmost edge indicates the end of the trial (which was always 8 seconds after the choice). Vertical lines, and changes in the bar texture, indicate Tcue and Treveal. The reward was always delivered 1.1 s after Treveal. (B) After the choice, an animated “hand of the clock” appeared on the left edge of the bar. The location of the hand indicated the current time during the trial. The hand moved rightward at a constant rate, eventually touching the right edge of the bar simultaneously with stimulus offset and the end of the trial. When the hand touched the first vertical line at Tcue, the line moved upward and morphed to become the cue. When the hand touched the second vertical line at Treveal, the line moved upward and revealed the outcome by ‘wiping out’ three of the possible reward bars, so that only the single reward bar that controlled the trial’s outcome remained on the screen.

**Figure S3.**
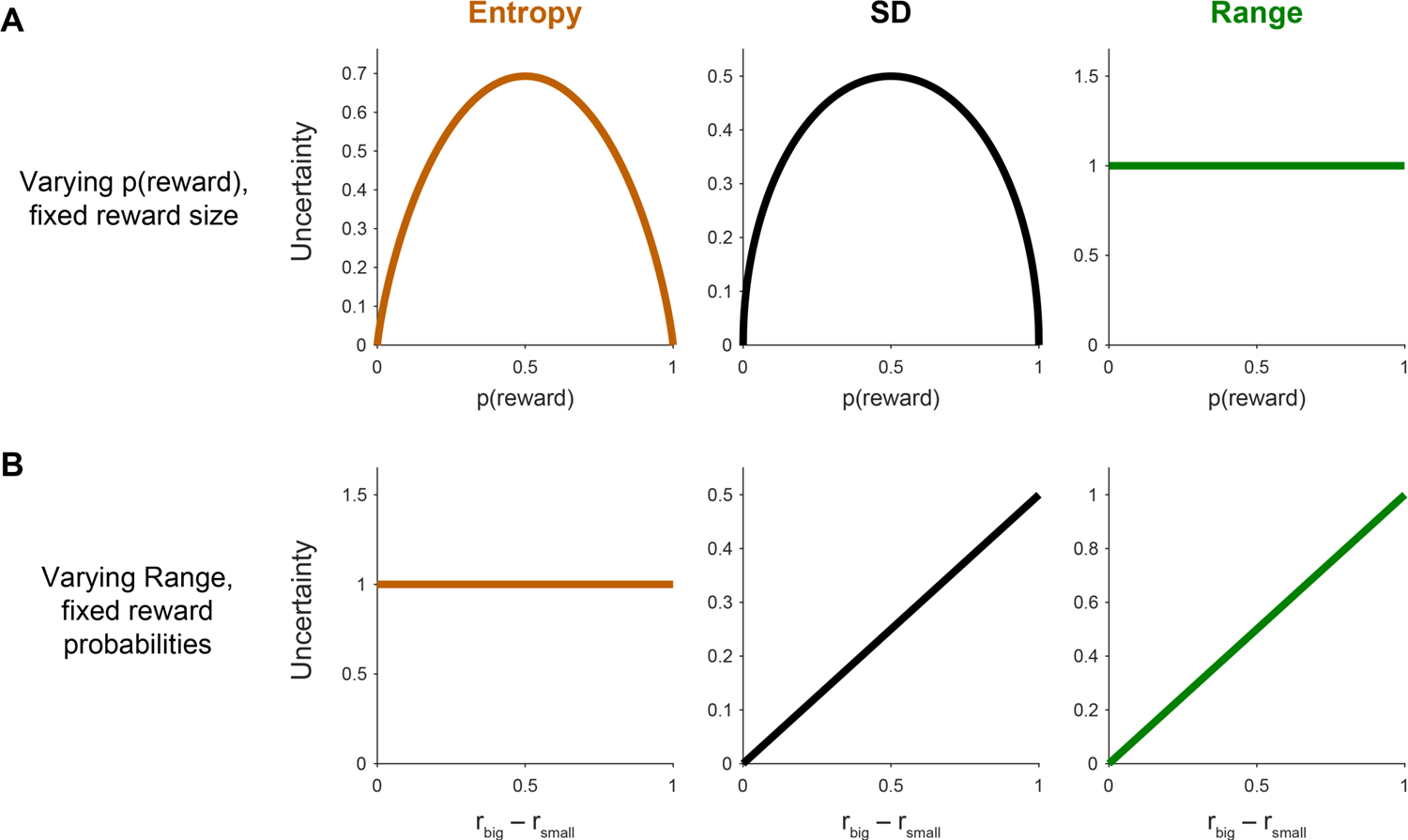
Necessity to manipulate multiple parameters of reward distributions to dissociate hypothesized forms of uncertainty. (A) One type of standard task used to study uncertainty has conditions with different reward probabilities while maintaining a fixed reward magnitude (e.g. (*65, 128, 129*)). This produces a classic inverted-U shaped function relating reward probability to certain measures of uncertainty (e.g. Entropy and SD; left and middle) but not other measures (e.g. Range; right), as shown in this plot in which the reward size is fixed at 1 and the reward probability is varied from 0 to 1. Thus, this type of task can measure behavior and neural activity related to certain forms of uncertainty, but cannot clearly distinguish between them, and cannot measure behavior and activity related to other forms of uncertainty (as has been pointed out by authors using those tasks (*129, 130*)). (B) A second type of standard task used to study uncertainty has conditions with different ranges between large vs small reward magnitudes while maintaining fixed reward probabilities (e.g. (*22, 50, 131*)). This produces a linear function relating reward range to certain measures of uncertainty (e.g. SD and Range; middle and right) but not other measures (e.g. Entropy; left), as shown in this plot in which the reward probabilities are fixed at p(big reward) = p(small reward) = 0.5 and the difference between the reward sizes is varied from 0 to 1. Thus, again, this type of task can measure behavior and activity related to certain forms of uncertainty, but cannot clearly distinguish between them, and cannot measure behavior and activity related to other forms of uncertainty (as has again been pointed out by authors using these tasks (*132*)).

**Figure S4.**
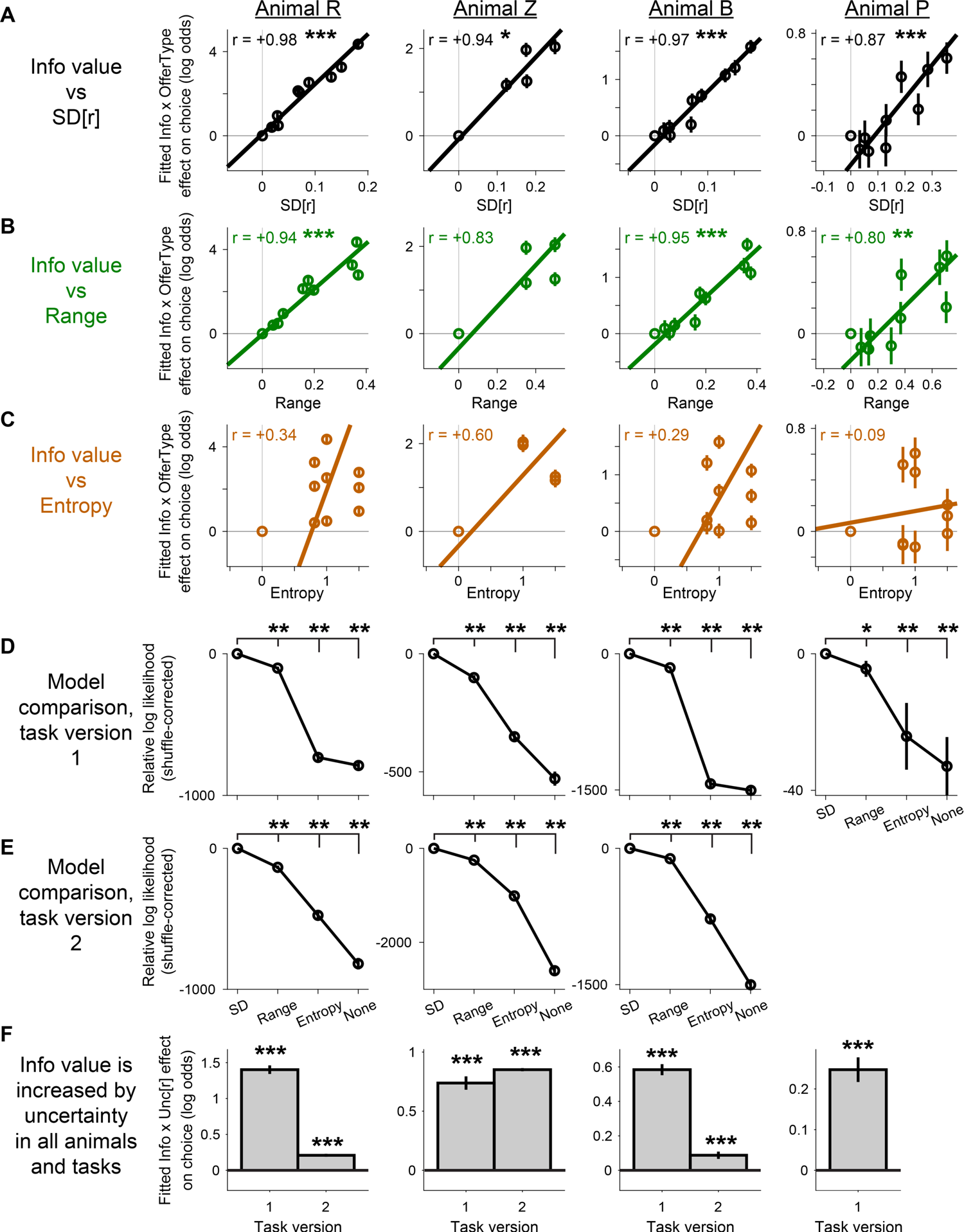
Consistent Info x Uncertainty Type effects across monkeys, tasks, and reward distributions. (A-C) Correlation between each candidate uncertainty measure – SD, Range, and Entropy, shown in panels A, B, and C – with the fitted Info x Uncertainty Type weights from a more detailed version of the model used in the main text, that included a more detailed classification of offers into uncertainty types (Table S2). In brief, this included all combinations of up to three types of reward distributions (25/50/25, 50/50, 25/75) x up to two types of risk levels (low range, high range), depending on which distributions were tested in each animal (Table S1). The weights are highly correlated with SD, less correlated with Range, and least correlated with Entropy. Error bars are SE. Text indicates Pearson’s linear correlation and asterisks indicate significance (*, **, *** indicate p < 0.05, 0.01, 0.001). The colored line is a linear fit from type 2 regression. (D) Direct comparison between models of behavior in which uncertainty is defined as SD, Range, or Entropy, or in which no uncertainty term is included (Table S2) separately for each animal, in the first version of the monkey task. Same format as Fig. 2H. The SD model fit best in all animals. (E) Same, in the second version of the monkey task, for all animals studied with the task. The SD model fit best in all animals. (F) Fitted weight of Info x Unc[r], measuring how the subjective value of information grows with uncertainty, from the main behavioral model for each animal and task (Table S1, model 1 (“Distrib SD”) for task version 1 and model 8 (“Clock SD”) for task version 2). All animals increased the value of information with uncertainty, as indicated by significantly positive weights. *, **, *** indicate p < 0.05, 0.01, 0.001.

**Figure S5.**
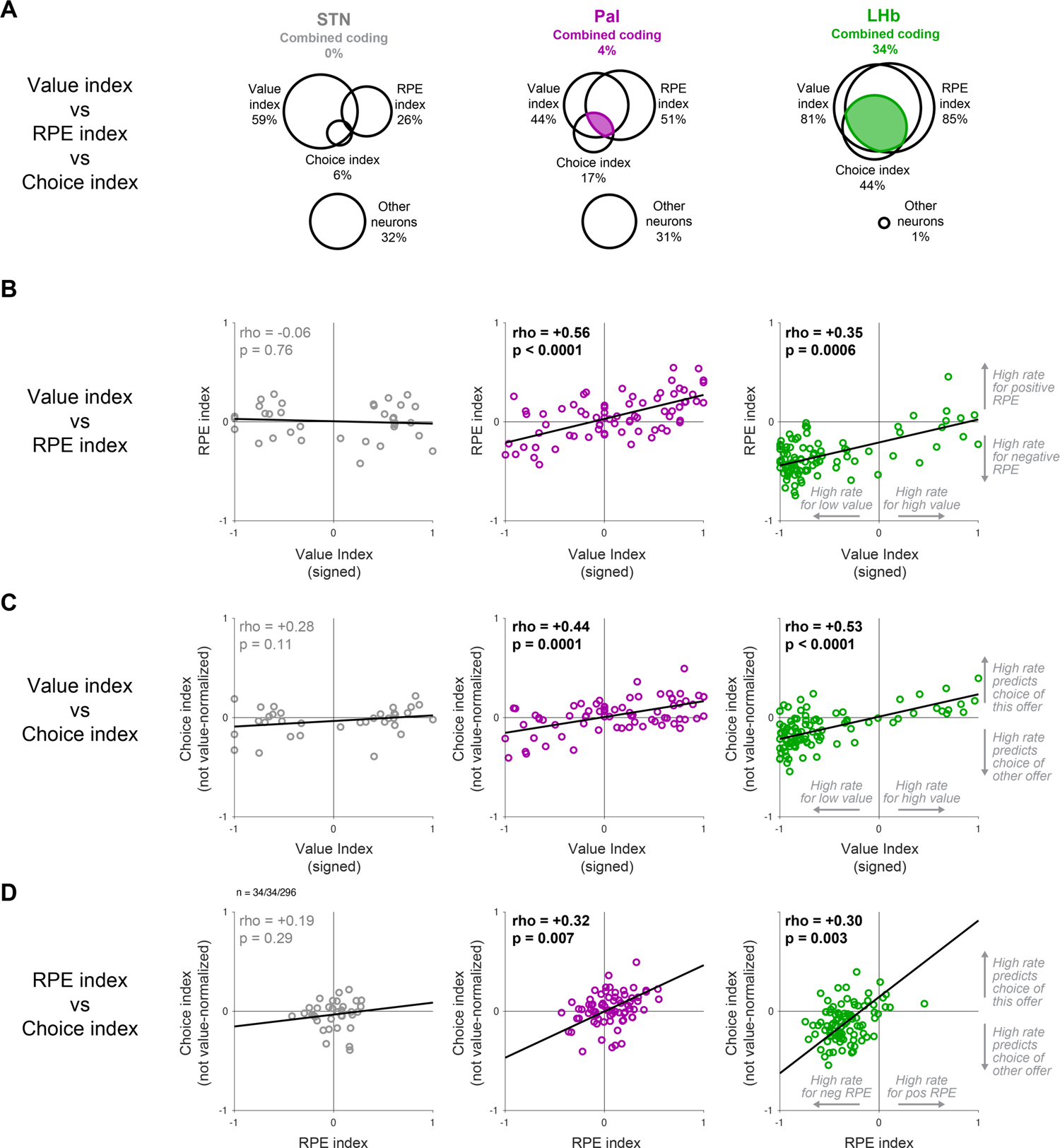
Value Coding, RPE Coding, and Choice Predictive Indexes are correlated in Pal and LHb, consistent with partial and full integration, but not STN. (A) Venn diagrams indicating overlap of the three coding indexes. Same as in Fig 4. Only LHb has a substantial population of neurons with a strong Value index, a significant RPE index, and significant Choice index, that all have the same coding sign. (B,C,D) Correlation of Value Coding Index with RPE Coding Index (B), Value Coding Index with Choice Predictive Index (C) and RPE Coding Index with Choice Predictive Index (D). All of these were highly significant in LHb and Pal, but not STN. This analysis was restricted to the subset of neurons for which all three indexes could be validly computed (n=34, 72, and 95 for STN, Pal, and LHb, respectively). Text indicates significance of rank correlation and its p-value. The line is the best linear fit by type 2 regression. For this analysis, to allow a direct comparison of the three indexes that can be interpreted in terms of changes in the neuron’s firing rate, all indexes were adjusted so that positive indexes indicate that higher firing rates were associated with higher offer values, more positive RPEs, and greater choice, respectively. Specifically, the Value Coding Index was multiplied by the sign of each neuron’s value coding effect (i.e. the sign of the mean of the fitted weight for the offer value term of the Value Models; models 17 and 18, Table S3), so that a positive value index in this plot means that higher firing rate was associated with higher offer value, while a negative value index means that higher firing rate was associated with lower offer value. Similarly, the Choice Predictive index for this analysis was computed as the simple correlation between residual normalized activity and residual choice (i.e. without converting residual normalized activity into a residual value signal by multiplying by the neuron’s sign of value coding, as was done in the main text), so that a positive index means that higher residual firing rate in response to an offer was associated with greater residual choice of the offer, while a negative index means that it was associated with lower residual choice of the offer. To incorporate data from both offers, the Choice Predictive Index for this analysis was computed by separately computing the indexes for offer 1 and offer 2 and then averaging the two.

**Figure S6.**
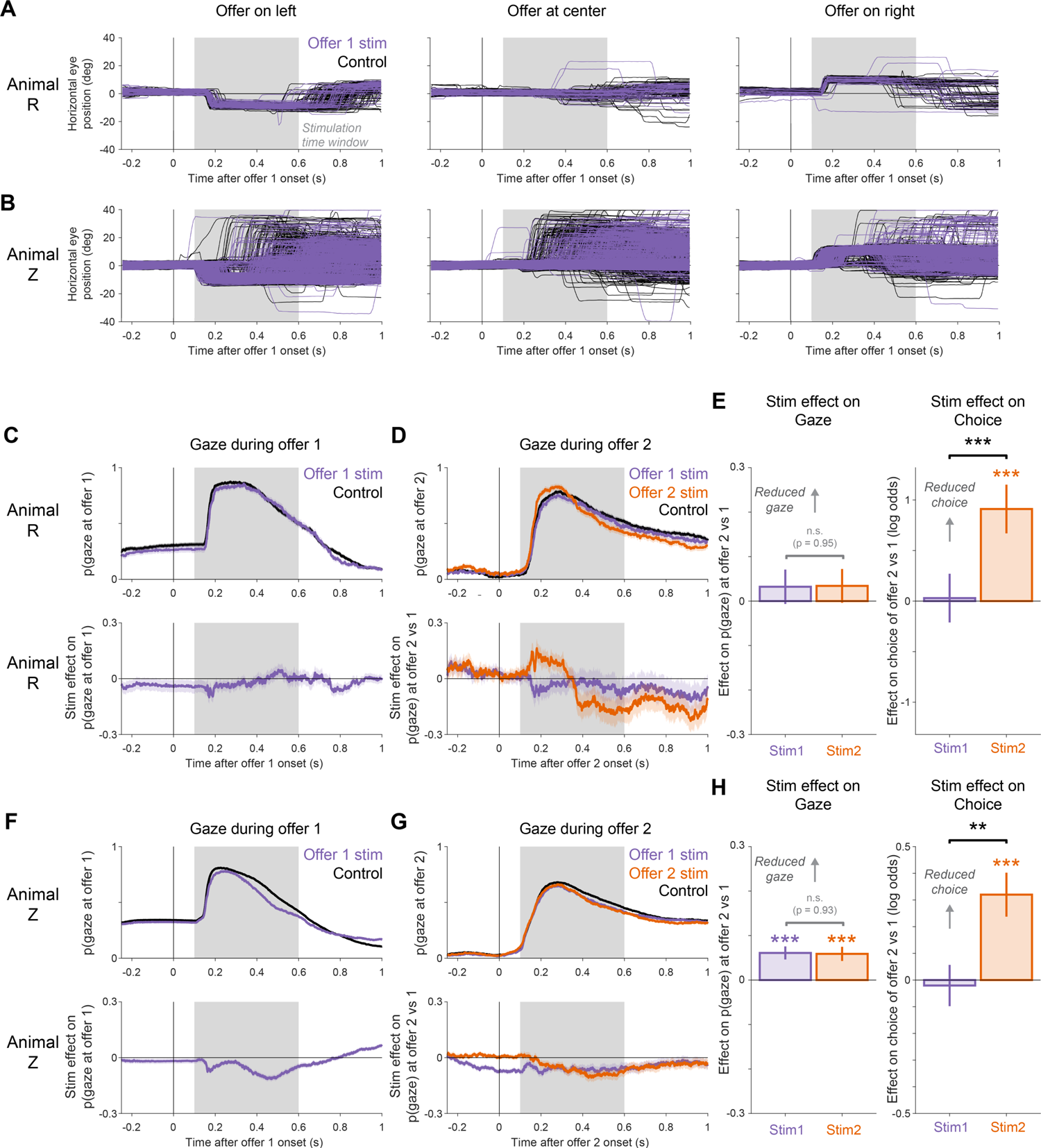
LHb stimulation does not produce motor commands to control gaze, and the causal effect of stimulation on choice cannot be explained by an effect of stimulation on gaze. (A-B) LHb stimulation does not produce motor commands that directly interfere with choice in our task. If stimulation did so, then it would produce stereotyped perturbations in gaze that are tightly time-locked to stimulation onset, as is commonly observed in oculomotor areas (e.g. superior colliculus and frontal eye fields (*133, 134*)). Furthermore, to interfere with choice, these perturbations would need to be both prominent and prolonged long after stimulation offset. For example, if stimulation induced contraversive eye movements, then it should induce a prominent rightward deviation in eye trajectories (since the electrode was located in the left LHb (Fig S7)). To test this, we compared eye position traces before, during, and after offer onset from single trials when the LHb was stimulated during offer presentation (purple) vs. a set of control trials in which no stimulation was performed during that offer (black; two control trials are shown for each stimulated trial). To make a direct comparison between full eye movement trajectories, this plot focuses on Offer 1 (to ensure the eye always began at the same position, i.e. the fixation point at the center of the screen before offer onset); shows the horizontal component of eye position (since the three possible offer locations were spaced horizontally on the screen); and shows all trials which produced continuous gaze trajectories (uninterrupted by blinks). Similar results were observed for both offers, for both horizontal and vertical eye positions, and when examining blinks. (A) Horizontal eye position during trials in which stimulation was performed during Offer 1 (purple curves) and control trials (black curves) in animal R, separately for offers presented on the left, middle, or right side of the screen (left, middle, right). Shaded area indicates the stimulation period. There was high overlap between the eye movement trajectories in stimulation and control conditions, including both initial saccades to the offer and later saccades away from the offer to empty locations on the screen. Stimulation did not induce obvious gross or stereotyped deviations from the natural variations of gaze shifts. (B) Same, for Animal Z. Again, there was high overlap between stimulation and control conditions. Stimulation did not induce obvious gross or stereotyped deviations from the natural variations of gaze shifts. (C-H) The causal effect of LHb stimulation on choice cannot be explained by an effect of stimulation on gaze. Specifically, while LHb stimulation did not have a direct motor-like effect on gaze, in some animals and conditions it had a modest tendency to influence the probability of gazing at the offer during and after stimulation, perhaps consistent with a change in the perceived motivational importance of the stimulus (C,D,F,G). Importantly, however, the effect on gaze alone could not explain the effect on choice: *choice* of Offer 2 vs Offer 1 was much more strongly influenced by stimulation during Offer 2, while *gaze* at Offer 2 vs Offer 1 was influenced similarly by stimulation during either offer (E,H). (C) Testing for an effect of Offer 1 stimulation on gaze during Offer 1 in animal R. *Top:* probability of gazing at Offer 1 (defined as gaze in a 6° x 7.5° rectangular window around the center of the offer) at each time before and after Offer 1 onset, separately for trials with Offer 1 stimulation (purple) and the remaining trials (black). Shaded area is ± 1 bootstrap SE. *Bottom:* estimated effect of Offer 1 stimulation on gaze at Offer 1 (purple), measured as the difference in gaze probability between the stimulation and control conditions above. There was little or no effect of stimulation on gaze during Offer 1. (D) Testing for effects of Offer 1 and Offer 2 stimulation on gaze during Offer 2 in animal R. *Top:* probability of gazing at Offer 2 at each time before and after Offer 2 onset, separately for trials with Offer 1 stimulation (purple), Offer 2 stimulation (orange), and the remaining trials (black). *Bottom:* estimated effect of stimulation on gaze bias between the two offers. We measured gaze bias as the difference in the probability of gazing at the two offers (p(gaze at Offer 2) – p(gaze at Offer 1)). We then estimated the stimulation effect as the difference between the gaze bias in each stimulation condition vs. the control condition, and quantified it as the mean effect in the stimulation window (gray shaded area). There were trends for Offer 1 stimulation to make gaze at Offer 2 slightly less likely, and for Offer 2 stimulation to make gaze at Offer 2 initially slightly more likely and later slightly less likely. However, neither effect reached significance (i.e. the bootstrap 95% CIs did not exclude 0). (E) Stimulation effects on gaze cannot explain stimulation effects on choice in animal R. *Left:* effect of stimulation during Offer 1 vs Offer 2 on gaze bias between Offer 1 vs Offer 2. This shows the same effects that were illustrated in (D), quantified here as the mean effect on p(gaze) during the stimulation window (gray shaded area), and plotted so that positive effects indicate a gaze bias against Offer 2. There were no significant effects of either Offer 1 or Offer 2 stimulation, and no significant difference between them (no bootstrap 95% CIs excluded 0). *Right:* effect of the same stimulations on the same trials on choice bias between Offer 1 vs Offer 2. This is the same plot as in Figure 5, quantifying choice bias using the fitted GLM in terms of the log odds of choosing Offer 1 vs Offer 2, plotted so that positive effects indicate a choice bias against Offer 2. There was no significant effect of Offer 1 stimulation, a highly significant effect of Offer 2 stimulation, and a highly significant difference between them. (F-H) Stimulation effects on gaze cannot explain stimulation effects on choice in animal Z. Same as (C-H), for animal Z. (F) There was a modest effect such that Offer 1 stimulation slightly reduced the probability of gazing at Offer 1 (bootstrap 99.9% CI excluded 0). (G) There was a modest effect such that either Offer 1 stimulation or Offer 2 stimulation slightly reduced the probability of gazing at Offer 2 (bootstrap 99.9% CIs excluded 0). (H) Stimulation effects on gaze were not significantly different between Offer 1 and Offer 2 stimulation (left), but stimulation effects on choice were significantly much greater for Offer 2 stimulation (right). *, **, *** indicate that the 95%, 99%, or 99.9% bootstrap CIs excluded 0 (left) or that the GLM regressor was significant at p < 0.05, 0.01, or 0.001, respectively.

**Figure S7.**
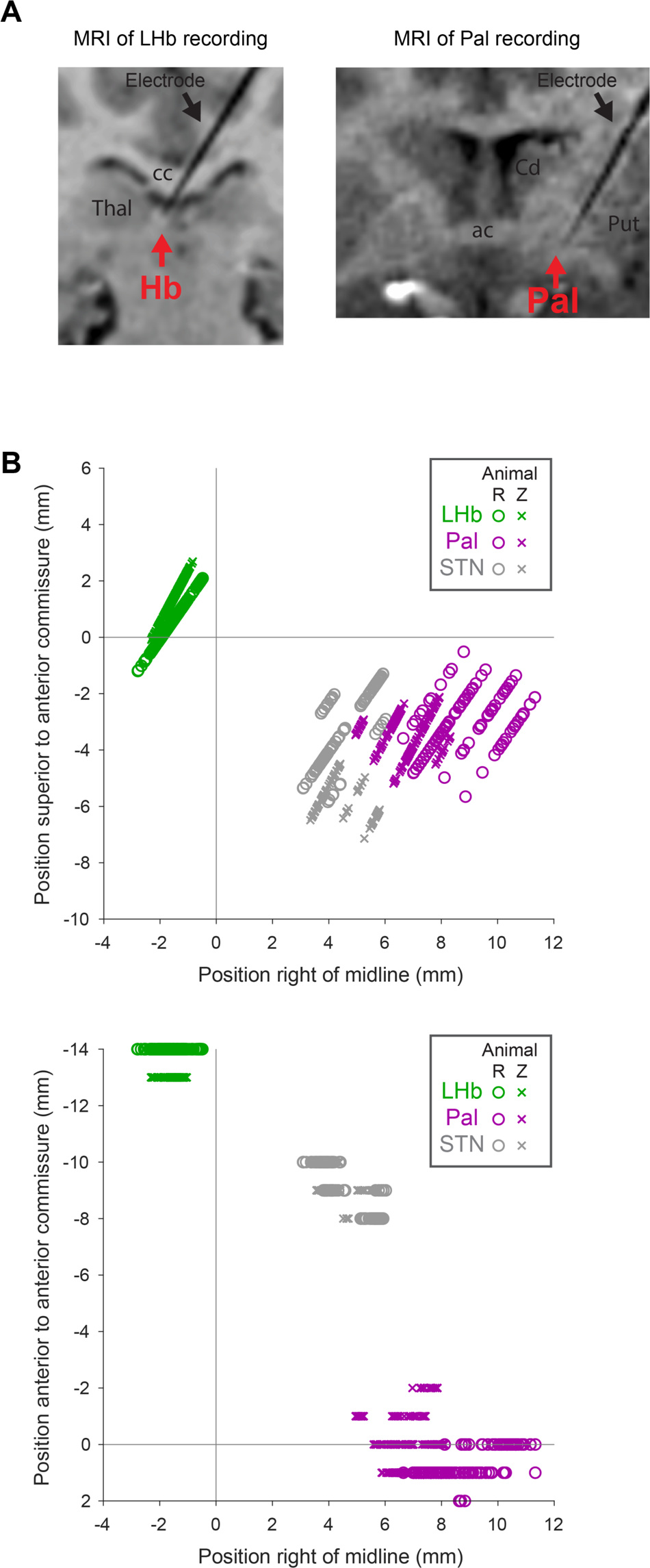
MRI verification and anatomical locations of recording sites. (A) MRIs taken immediately after recording with the electrode still in place, verifying its location in the target area. Shown are coronal views slightly tilted to align with the electrode track. The electrode is visible as a black ‘shadow’ on the MRI. Black arrows indicate the electrode. Red arrows indicate the target area. Left: a recording site in LHb. Right: a recording site in Pal. Abbreviations: Hb, habenula; Pal, pallidum; cc, corpus callosum; Thal, thalamus; ac, anterior commissure; Cd, caudate nucleus; Put, putamen. (B) Anatomical locations of recording sites. Reconstructed 3D coordinates of each neuron in the dataset, shown for all areas (indicated by colors: LHb, Pal, and STN as green, purple, and gray, respectively) and animals (indicated by symbols: circles and crosses for animals R and Z, respectively). Coordinates are relative to the midline, superior tip of the anterior commissure. *Top* shows coordinates in the coronal plane. *Bottom* shows coordinates in the horizontal plane. The coordinates for each area were anatomically distinct from each other and were similar in both animals.

**Table S1.** Offer reward distribution statistics for all animals and sessions used in behavioral analysis. Each row shows the parameter settings used to generate the offers while collecting a behavioral dataset after animals had been trained at the task and that was used in the main analysis of behavior. Separate data was used for analysis of behavior under normal conditions (tasks “1” and “2”) and in sessions with stimulation (task “2 w/stim”). Animal B’s behavioral dataset from task 2 included data collected with three slightly different task parameters as indicated by separate rows in the table; each other dataset from a given animal and task condition used a single set of parameters. All reward amounts and ranges are in units of mL juice. Columns indicate the animal’s identity; the task version (version 1 where offers showed reward distributions, or version 2 where offers also showed a clock); the number of correctly performed trials collected (only counting trials of the multi-attribute information task, not the interleaved simple information anticipation task if it was present in those sessions); the possible expected reward values for offer 1, indicated as a min-max range; the possible expected reward values for offer 2, indicated either as a min-max range (meaning that it was drawn independently of offer 1) or as “+/-“ (indicating that it was drawn from a range relative to the expected reward value of offer 1); the possible reward ranges for uncertain reward distributions, indicated either as a min-max range of possible reward ranges, or a set of specific reward ranges; the probability of the offer having each of the possible reward distribution types (Safe, 25/50/25, 50/50, or 25/75); and the settings of Rstep and Rmax.

| Animal | Task | #trials | E[r] off 1 | E[r] off 2 | Range | P(reward distribution type) |  |  |  | Rstep | Rmax |
| --- | --- | --- | --- | --- | --- | --- | --- | --- | --- | --- | --- |
|  |  |  |  |  |  | Safe | 25/50/25 | 50/50 | 25/75 |  |  |
| R | 1 | 10295 | 0.14-0.46 | +/- 0.24 | 0-0.6 | 0.11 | 0.33 | 0.33 | 0.22 | 0.02 | 0.6 |
| Z | 1 | 6159 | 0.15-0.55 | 0.15-0.55 | (0.35,0.5) | 0.33 | 0.33 | 0.33 |  | 0.025 | 0.7 |
| B | 1 | 21038 | 0.14-0.46 | +/- 0.24 | 0-0.6 | 0.11 | 0.33 | 0.33 | 0.22 | 0.02 | 0.6 |
| P | 1 | 15116 | 0.28-0.85 | +/- 0.32 | 0-1.1 | 0.11 | 0.33 | 0.33 | 0.22 | 0.0375 | 1.125 |
| R | 2 | 40434 | 0.42-0.78 | 0.42-0.78 | (0.6,0.84) | 0.33 | 0.33 | 0.33 |  | 0.04 | 1.2 |
| Z | 2 | 87491 | 0.48-0.72 | +/- 0.1 | (0.6,0.84) | 0.33 | 0.33 | 0.33 |  | 0.04 | 1.2 |
| B | 2 | 4352 | 0.28-0.92 | +/- 0.5 | 0.3-1.2 | 0.33 | 0.22 | 0.22 | 0.22 | 0.04 | 1.2 |
| B | 2 | 16667 | 0.42-0.78 | +/- 0.35 | 0.4-1.2 | 0.33 | 0.22 | 0.22 | 0.22 | 0.04 | 1.2 |
| B | 2 | 13870 | 0.42-0.78 | 0.42-0.78 | (0.84) | 0.33 | 0.33 | 0.33 |  | 0.04 | 1.2 |
| R | 2 w/stim | 1016 | 1.05-1.95 | 1.05-1.95 | (1.5,2.1) | 0.33 | 0.33 | 0.33 |  | 0.10 | 3.0 |
| Z | 2 w/stim | 7822 | 0.48-0.72 | +/- 0.1 | (0.6,0.84) | 0.33 | 0.33 | 0.33 |  | 0.04 | 1.2 |

**Table S2.**
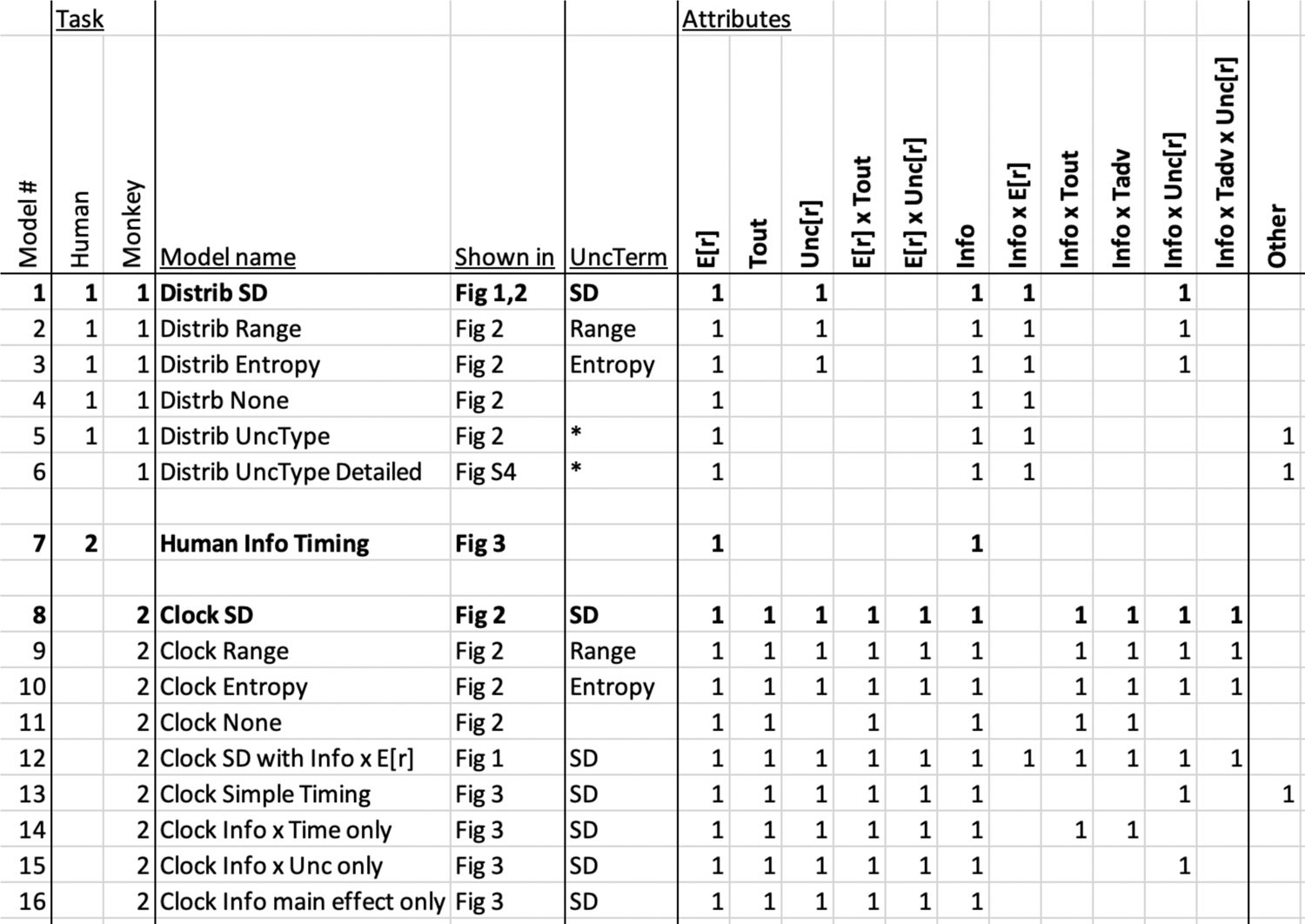
Models used to fit behavior and/or neural activity to examine the attributes underlying the subjective value of information and reward. Each row is one generalized linear model. Bold rows are the main models used for modeling the behavioral or neuronal data collected from each human or monkey task (“Distrib SD”, “Human Info Timing”, and “Clock SD”). Other rows are alternate models used for model comparisons or to analyze the influence of specific attributes. Columns indicate the model number; the human and monkey task versions whose data the models were used to fit; the model name; the figures where it was used; the uncertainty term (if any) used in the model, with * indicating models that did not use only a single term for uncertainty and instead had separate terms for separate types of uncertain reward distributions (as described in Methods); which offer attributes were included in the model (attributes marked with a “1” were included, and otherwise were not included); and whether other additional attributes were included. Specifically, as described in Methods, the “Distrib UncType” and “Distrib UncType Detailed” models had separate attributes for whether an offer had each of several specific types of uncertain reward distributions, as well as the interactions of those uncertainty types with Info; similarly, the “Clock Simple Timing” model had separate weights for whether an offer had each of several specific settings of (Tcue, Treveal) event timings. Finally, in addition to the attributes listed here, all neural and behavioral models included the appropriate set of bias terms, e.g. fits to neural activity included a constant term representing the mean or baseline firing rate (Methods).

**Table S3.** Models used to fit behavior and/or neural activity to examine the neural implementation of total subjective value. Each row is one generalized linear model. Columns indicate the model number; the monkey task version whose data the models were used to fit; the model name; the figures where it was used; the attributes used in the model; and whether other attributes were used. For all of these models, the values of the two offers (V1 and V2) were derived from fits of the main model for the second version of the monkey task (“Clock SD”; Methods). For each LHb neuron, LHbResid1 and LHbResid2 were derived from the residuals from fitting that model to the neuron’s responses to Offer1 and Offer 2, respectively (Methods). For each stimulation session, Stim1 and Stim2 indicated whether electrical stimulation was delivered during Offer1 and Offer 2, respectively. Finally, “StimPred” models were used in the model comparison to determine the attributes which interacted with stimulation effects. The “StimPred Stim2xAttrib” model is marked as “other” because it is the result of a model selection procedure which iteratively adds parameters to the model based on whether they significantly improve the fit (Methods). Finally, in addition to the attributes listed here, all neural and behavioral models included the appropriate set of bias terms, e.g. fits to neural activity included a constant term representing the mean or baseline firing rate (Methods).

| Model # | Monkey Task | Stimulation | Model name | Shown in | Attributes |  |  |  |  |  |  |  |  |  |  | Other |
| --- | --- | --- | --- | --- | --- | --- | --- | --- | --- | --- | --- | --- | --- | --- | --- | --- |
|  |  |  |  |  | V2-1 | V1 | V2 | LHbResid2-1 | LHbResid1 | LHbResid2 | Stim2-1 | Stim1 | Stim2 | Stim2 x (V2-1) | Stim2 x V2 |  |
| 17 | 2 |  | "Value Model" for Offer 1 | Fig 4 |  | 1 |  |  |  |  |  |  |  |  |  |  |
| 18 | 2 |  | "Value Model" for Offer 2 | Fig 4 |  |  | 1 |  |  |  |  |  |  |  |  |  |
| 19 | 2 |  | ChoicePred | Fig 4,5 | 1 |  |  | 1 |  |  |  |  |  |  |  |  |
| 20 | 2 |  | ChoicePred by Offer | Fig 4,5 | 1 |  |  |  | 1 | 1 |  |  |  |  |  |  |
| 21 | 2 | 1 | StimPred by Offer | Fig 5 | 1 |  |  |  |  |  |  | 1 | 1 |  |  |  |
| 22 | 2 | 1 | StimPred None | Table S4 | 1 |  |  |  |  |  |  |  |  |  |  |  |
| 23 | 2 | 1 | StimPred Stim2 | Table S4 | 1 |  |  |  |  |  |  |  | 1 |  |  |  |
| 24 | 2 | 1 | StimPred Stim2xV2-1 | Table S4 | 1 |  |  |  |  |  |  |  | 1 | 1 |  |  |
| 25 | 2 | 1 | StimPred Stim2xV2 | Table S4 | 1 |  |  |  |  |  |  |  | 1 |  | 1 |  |
| 26 | 2 | 1 | StimPred Stim2xAttrib | Table S4 | 1 |  |  |  |  |  |  |  | 1 |  |  | 1 |

**Table S4.**
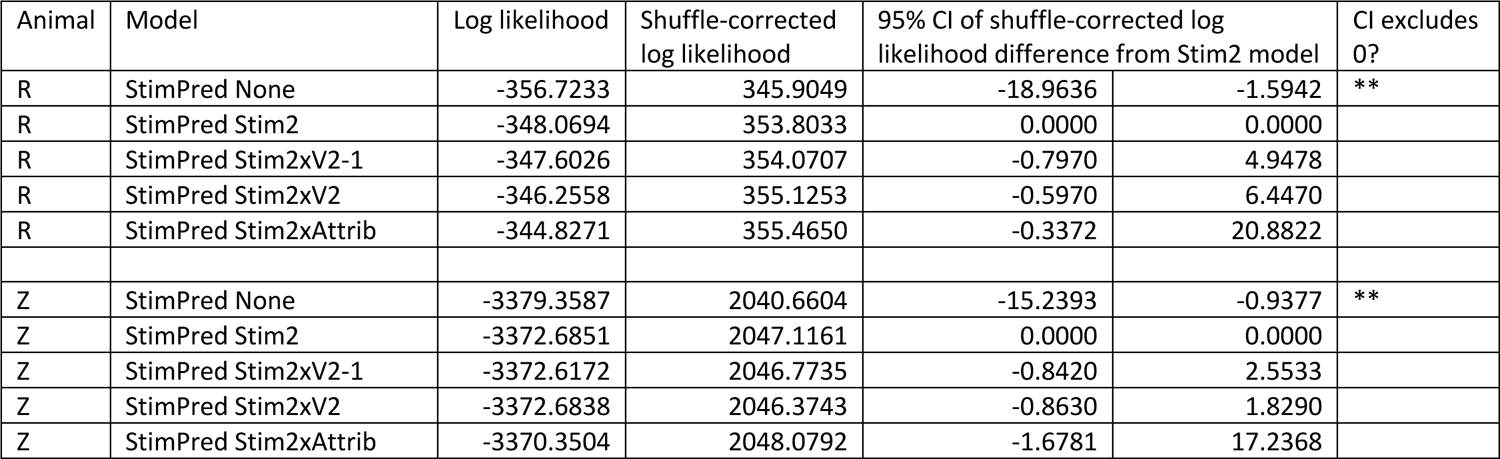
The influence of LHb offer 2 stimulation on choice was predominantly explained by a main effect of stimulation. Each row is one model fit (see Methods and Table S3 for full descriptions of all models). In brief, the “StimPred None” model does not include any stimulation effects; the “StimPred Stim2” model includes a main effect of stimulation during offer 2; and the further models additionally include interactions between stimulation and the value difference between the offers (“Stim2xV2-1”), the value of offer 2 (“Stim2xV2”), or a subset of offer 2 attributes selected via a stepwise model selection procedure to optimize the fit (“Stim2xAttrib”; Methods). Columns indicate the animal, model, log likelihood, shuffle-corrected log likelihood, and the 95% bootstrap confidence interval of the difference between the shuffle-corrected log likelihoods of the current model and the “StimPred Stim2” model. *, **, *** indicate that 0 was excluded by the 95%, 99%, or 99.9% bootstrap confidence intervals. In both animals, the fit was reliably improved by including the main effect of Stim2 (“StimPred Stim2” vs “StimPred None”), but was not reliably improved by including interactions between Stim2 and either offer values or specific offer attributes.

| Animal | Model | Log likelihood | Shuffle-corrected log likelihood | 95% CI of shuffle-corrected log likelihood difference from Stim2 model |  | CI excludes 0? |
| --- | --- | --- | --- | --- | --- | --- |
| R | StimPred None | -356.7233 | 345.9049 | -18.9636 | -1.5942 | ** |
| R | StimPred Stim2 | -348.0694 | 353.8033 | 0.0000 | 0.0000 |  |
| R | StimPred Stim2xV2-1 | -347.6026 | 354.0707 | -0.7970 | 4.9478 |  |
| R | StimPred Stim2xV2 | -346.2558 | 355.1253 | -0.5970 | 6.4470 |  |
| R | StimPred Stim2xAttrib | -344.8271 | 355.4650 | -0.3372 | 20.8822 |  |
| Z | StimPred None | -3379.3587 | 2040.6604 | -15.2393 | -0.9377 | ** |
| Z | StimPred Stim2 | -3372.6851 | 2047.1161 | 0.0000 | 0.0000 |  |
| Z | StimPred Stim2xV2-1 | -3372.6172 | 2046.7735 | -0.8420 | 2.5533 |  |
| Z | StimPred Stim2xV2 | -3372.6838 | 2046.3743 | -0.8630 | 1.8290 |  |
| Z | StimPred Stim2xAttrib | -3370.3504 | 2048.0792 | -1.6781 | 17.2368 |  |

## Supplemental Discussion

### The LHb and online decision making

We demonstrate converging evidence that LHb neurons reflect value and influence online decisions: (1) the lion’s share of LHb offer responses were explained by subjective value; (2) variations in these value signals were choice-predictive; (3) LHb stimulation altered decisions as if the value of the stimulated offer was reduced; (4) both the choice-predictiveness of LHb value signals and the causal influence of LHb stimulation were greater when both offers were presented and animals could decide between them. Thus, the LHb does not simply signal RPEs for the purpose of adjusting estimated values over long timescales during gradual reinforcement learning (e.g. across multiple trials or presentations); these signals can also directly influence ongoing multi attribute decisions.

This finding may explain why LHb neuronal activity has been linked to the speed of actions (*135, 136*) and blood-level oxygenation dependent signals have been linked to the willingness to initiate actions (*104*). Indeed, the LHb has suitable projections to implement this influence, which could be mediated by its strong control over midbrain circuitry that regulates neuromodulators (*137*) including dopamine neurons (*138*), whose projections to basal ganglia circuits have been implicated in both learning and immediate approach or withdrawal from motivational goals (*139*), and the serotonin system (*140*), which has been implicated in learning, motivational, and decision-making deficits and associated with psychiatric disorders (*141–145*).

An important goal of future work will be to uncover the mechanisms by which LHb activity influences decisions in different environments. We found that LHb integrates many attributes of both information and reward into its value signal, including: expected reward, reward uncertainty, reward timing, the opportunity to gain information, the degree to which information value scales with reward uncertainty, and the degree to which information value scales with the timing of information in advance of the reward. These were all integrated together in a manner closely resembling the subjective value that drives decisions. However, even the high-dimensional space of offer attributes we probed with this task is only a subspace of the much larger space of attributes that organisms can consider when making decisions in natural environments. Hence future work will be needed to assess whether and how the LHb integrates other distinct types of attributes into its subjective value-related signals during decisions, such as novelty (*124*), aversion (*73*), aggression (*146*), and drug reward (*147*).

Our data implicate the LHb in a type of decision making that is ubiquitous in natural environments – decisions between complex, multi-attribute options with uncertain reward outcomes, where individuals can evaluate each option as it arises and have time to deliberate before making a choice. Under these conditions, our data indicates that LHb activity tracks subjective value and both predicts and influences the decision. However, it is possible that the LHb has a different degree of involvement in other types of decision making. For example, there is evidence that LHb value-related responses may be too slow to have an online influence on very rapid or automatic decisions (e.g. immediate saccades), and in such cases may primarily signal RPEs after the decision has been made (*35, 148*). LHb may also have less influence on simple single attribute decisions (e.g. big vs small reward (*76, 100*)), and would require additional signals and processing to be ‘bound’ to specific offers if multiple offers are presented simultaneously (*149*). Hence future work will be needed to determine whether and how the LHb influences other types of decision making.

### Information seeking and multi-attribute decision making

An important advance necessary to demonstrate our behavioral and neuronal findings was our task design allowing us to probe the detailed structure of uncertainty and value judgements during multi-attribute decision making, and to identify neurons reflecting these computations. This was critical to draw accurate conclusions about value computations.

Notably, we found that a large number of neurons in both Pal and LHb were sensitive to multiple reward- and/or information-related attributes. Neurons in both areas responded significantly to similar numbers of offer attributes, and produced at least partially integrated value-like signals (Fig 4). Furthermore, neurons in both areas encoded multiple motivational signals in correlated manners, including offer value, choice-predictive activity, and RPEs (Fig S5). Thus, if we had used a simpler task that manipulated information and reward in simple binary manners (e.g. only manipulating information through a binary Info vs Noinfo distinction, and only manipulating primary reward through a binary big vs. small reward distinction; without manipulating uncertainty or time), then it might have been tempting to interpret the resulting data to conclude that many neurons in both Pal and LHb integrated reward and information and were consistent with tracking subjective value.

Instead, we were able to make a much more precise distinction between partial and full integration, by manipulating additional motivational attributes like uncertainty and timing (that are both valuable in their own right and also interact with information and primary reward to alter their values), and then modeling the precise manner in which individuals integrated them in the value judgements underlying their decisions. This allowed us to conclude that many LHb neurons fully integrated attributes consistent with subjective value, while this was much rarer in Pal (Fig 4); and that many of these LHb neurons further combined their offer value signal with two other coding properties, choice-predictive activity and RPEs, a combination which was very rare in Pal (Fig S5).

This multi-attribute nature of the task design, and the resulting conclusions it enabled, are especially crucial given that most real-life decisions occur in situations where both information and extrinsic outcomes have multiple attributes that can be traded off against each other, including expected reward, uncertainty, and timing. For example, a concealed hunter stalking its prey must decide between taking a peek to gather information (at the risk of alerting the prey), or remaining under cover (at the risk of losing the prey or taking longer to track it down).

### Information seeking, reward seeking, and uncertainty

There has long been intense investigation into how organisms adjust their behavior to handle uncertain environments. Yet there is still considerable debate over the fundamental question: *what form* of uncertainty motivates behavior?

Indeed, there is even debate over whether many such uncertainty-related behaviors involve any internal estimate of uncertainty at all. For example, economic theories of risky decision making have long included proposals that individuals explicitly compute a measure of risk, such as the variance of the outcome distribution, and directly place value on it (e.g. mean-variance theory (*64*)). However other, foundational economic theories long proposed that individuals need not explicitly compute a measure of risk and instead behave as if they use a nonlinear utility function over outcomes, such that an outcome distribution with greater dispersion will tend to be assigned lower expected utility (concave utility function) or greater expected utility (convex utility function) (e.g. expected utility theory (*150*)). An analogous debate has taken place about information-seeking behavior, ever since the first discovery of ‘observing behavior’ in the 1950s, with different psychological and economic theories proposing that individuals value information either explicitly, based on an internal measure of uncertainty, or implicitly, as a side-effect of a nonlinear conditioned reinforcement function ((*11, 17, 23, 44-46, 151, 152*); for review see (*7*)).

Furthermore, if a form of behavior is motivated by an internal estimate of uncertainty, then understanding that behavior fundamentally requires understanding the functional form the brain uses to measure that uncertainty, and the neural circuitry implementing that computation. Theories of risk-related decision making founded in economics have often focused on the variance, which approximates risk attitudes arising from utility functions in certain situations (*153*). Neuronal recordings in economic decision tasks have modeled risk-related activity as functions of variance or standard deviation, or using nonlinear utility functions from economic theory (*50, 131, 154-156*). However, many studies have reported another form of potentially uncertainty-related activity, in which neural activity during sensory processing or economic decision tasks can rescale with the dispersion of possible outcomes, and this rescaling has often been modeled as a function of standard deviation (*157, 158*) or range (*60, 63, 132*). By contrast, work on information seeking has proposed a gamut of uncertainty measures that could motivate information seeking, including entropy (*44, 53, 159*), standard deviation or variance (*13, 14*), intermediates between the two (*37, 160*), and implicit functions of state values and prediction errors (*11, 46, 68*).

In order to distinguish these possibilities to determine what putative uncertainty measure the brain uses to compute the value of information about future rewards, we designed a task in which individuals could be directly shown rich probability distributions of reward outcomes, allowing us to manipulate uncertainty by varying both the possible reward amounts and their probabilities. This multi-dimensional manipulation of uncertainty is necessary to dissociate families of uncertainty measures. Intuitively, if uncertainty is manipulated using only a single parameter of the reward distribution, then the results could always be consistent with many possible uncertainty measures (or families of uncertainty measures), as long as those measures are all correlated with that parameter. For example, the common manipulation of uncertainty by manipulating only reward probability cannot distinguish between SD and entropy, while manipulating only reward range cannot distinguish between SD and range (Fig S3). Hence we addressed this by developing a task where individuals are offered full probability distributions of outcomes, allowing a direct test between these families of uncertainty measures.

We found that humans and monkeys scaled the value of information with an uncertainty measure resembling the standard deviation or variance of rewards (Fig 2, S4). Furthermore, Pal and LHb neurons scaled their information signals in a highly similar manner, consistent with a role in this value computation. This provides crucial evidence that the value of information tracks the suite of reward statistics that produce uncertainty in natural environments, including both probabilities and magnitudes, and places important constraints on the underlying neural computations.

In particular, our data implies that the partially and fully integrated information signals present in Pal and LHb must have been originally computed by brain areas with access to specific offer attributes, such as reward probabilities and reward amounts of the multiple potential outcomes of each offer. This is necessary to compute SD-like uncertainty measures, because they require knowledge of both the probabilities and amounts of all possible outcomes. This sophisticated knowledge about reward distributions could potentially be encoded through one of several proposed neural representations of probability distributions, including subsets of neurons that explicitly encode specific attributes of distributions like reward probabilities (*69*) or population-based distributional codes (*161–163*)). Furthermore, our data implies that after being computed, this full distributional information must then be distilled down to a single scalar measure of ‘total amount of uncertainty’, through a function resembling the SD, which is then used to scale Pal and LHb information-related signals and the subjective value of information that guides decisions (Fig 2).

By contrast, other families of uncertainty measures would be better suited for other functional roles in the brain. Measures like entropy only track probabilities, so they may be most useful in situations where the possible ‘outcomes’ can have different probabilities but are all of similar motivational importance (e.g. sensory areas concerned with the statistics of natural scenes, e.g. the motion, orientation, and form of visual objects (*55*)). Measures like range only track extreme magnitudes, so they may be most useful for setting the dynamic range of neural responses (e.g. (*62*)). An important question for future work will be to discover what forms of uncertainty guide information seeking in other contexts, such when outcomes are aversive (*24, 164*), when offers are ambiguous or have hidden attributes (*165*), and when information is instrumental (*165–168*).

Finally, these findings also provide crucial support for the class of theories of information seeking described at the start of this section, which propose that information is valued based on an explicit internal estimate of the amount of uncertainty it will resolve and the time it will resolve it. We find that many LHb neurons have activity related to the total, integrated subjective value of an offer. In principle, one could imagine that these subjective values could have been computed using an algorithm that accounts for uncertainty either explicitly (e.g. using a direct estimate of the ‘amount of uncertainty’ (*23, 44*)) or implicitly (e.g. via a nonlinear form of conditioned reinforcement (*8, 45, 46*)). However, crucially, we also find many Pal neurons that do not necessarily encode total subjective value, but *do* encode an offer’s informativeness, and the interactions of informativeness with uncertainty, time, or both (Fig 4, S4). Thus, we show that neurons in a nucleus with strong projections to LHb have activity resembling explicit representations of the amount of uncertainty and of the time that uncertainty will be resolved by information. These representations closely resemble the influences of these attributes on value (Fig 1-3), emerge immediately after offer presentation in the same epoch when LHb activity emerges reflecting the total subjective value of offers in decisions (Fig 4) and the LHb has a causal influence on behavior (Fig 5). Thus, our data shows that key basal ganglia circuitry contains explicit representations of uncertainty and future opportunities to gain information, in the right time and place to contribute to computing the value of information and to motivating information seeking decisions.

